# Soybean *Rhg1* resistance to *Heterodera glycines* hinders the parasitism of *Pratylenchus penetrans*

**DOI:** 10.1101/2025.09.29.679389

**Authors:** Deepak Haarith, Ann E. MacGuidwin, Andrew F. Bent

## Abstract

*Rhg1* has been the most effective QTL deployed in soybeans (*Glycine max*) to control soybean cyst nematode (*Heterodera glycines*; SCN). However, the resistance mechanisms and specificity of *Rhg1* towards SCN as opposed to other plant parasitic nematodes are not well known. In this study we report that *Rhg1* can <u>hinder</u> the parasitism of the root lesion nematode *Pratylenchus penetrans* (Pp), which is commonly found along with SCN in northern U.S. soybean fields. Two elite public soybean varieties, carrying either *rhg1-a* + *Rhg4* or *rhg1-b*, negatively impacted Pp populations measured at 30 dpi. This contribution of *Rhg1* was more rigorously demonstrated using near isogenic lines for *rhg1-b*. Additionally, in three of the four soybean genetic backgrounds tested we observed significantly more Pp at 30 dpi when Pp were co-inoculated with SCN, consistent with a previous study. No significant changes in SCN cyst numbers upon co-inoculation with Pp were observed for any of the SCN-resistant *rhg1* lines. No nematode-induced changes in *Rhg1* transcript abundances were observed when sampling infected root sections in any of the plant genotypes, and no changes in a jasmonate response indicator were observed. However, the 3 dpi salicylate-mediated response to SCN or Pp became more robust during SCN + Pp co-infection. Multiple hypotheses for further dissection and manipulation of soybean resistance to *Pratylenchus* root lesion nematodes are suggested based on the present findings.

## Introduction

Root lesion nematodes (RLN^a^; *Pratylenchus* spp.) are important agricultural pests (Jones et al. 2013; Rosa de Araújo et al. 2025). They are migratory ectoparasitic and endoparasitic nematodes (Zunke 1990) with a wide host range including soybean.

*Pratylenchus brachyurus* is often considered to be the most damaging nematode pest of soybeans in South America (Castanheira et al. 2020; Favoreto et al. 2019; Nomura et al. 2024). *Pratylenchus* populations survive cold winters in dead roots, including through anhydrobiosis or slow desiccation (Townshend 1984). *Pratylenchus* are some of the most common plant parasitic nematodes found in the soybean producing midwestern U.S. states like Wisconsin, Minnesota, Iowa, and Illinois, with *Pratylenchus penetrans* (Pp) being among the most common species, and a species for which soybean yield losses have been documented (Chen et al. 2012; MacGuidwin and Bender 2012; MacGuidwin et al. 2023; Saikai and MacGuidwin 2022; Schmitt and Barker 1981; Schumacher and Bird 2025). Unlike sedentary endoparasitic nematodes such as the soybean cyst nematode (SCN, *Heterodera glycines*), Pp feed on soybean roots at multiple sites as ectoparasites or endoparasites (Endo 1967; Saikai and MacGuidwin 2020). SCN spend limited time exposed to environmental disturbances as they shelter inside host roots or in cyst-encased eggs for much of their life cycle. In contrast, the migratory lifestyle of Pp (Zunke 1990) may make them more directly exposed to, and to some extent adapted to, changes in soil pH and nutrients, soil moisture, soil type, own population density, microbial communities, and competition from other nematodes. SCN is the most economically important pathogen or pest on soybeans in the United States (Bradley et al. 2021). Across all of the North Central U.S. and Ontario for ten recent years, SCN is estimated to have caused approximately 4% soybean yield loss (about 1/4 of the total for all soybean pathogens) (Allen et al. 2017; Bradley et al. 2021). Understandably, most efforts in U.S. soybean systems have been directed at managing SCN and not *Pratylenchus* spp. Estimates of the impact of RLN on soybean production are rare, but RLN and SCN frequently co-occur (MacGuidwin et al. 2023; Saikai and MacGuidwin 2022; Schumacher and Bird 2025). Northern U.S. soybean fields are typically rotated with maize, an excellent host for RLN. In one deeply studied set of Wisconsin soybean fields, RLN was found to cause up to 4.5% yield loss by itself (Saikai and MacGuidwin 2022), and about 3.79% yield loss on corn (MacGuidwin and Bender 2016). In 2024 the farm gate values of corn and soybean in Wisconsin alone were over USD $2.26 billion and USD $1.01 billion respectively (https://www.nass.usda.gov/Quick_Stats/Ag_Overview/stateOverview.php?state=WISCON SIN), which illustrates the economic impact of even a 1% yield loss to RLN.

Along with crop rotation, the most important tool for managing SCN over the past three decades has been host resistance provided by the major-effect quantitative trait locus (QTL) *Rhg1* (Resistance to *Heterodera glycines* 1) (Bent 2022; Niblack et al. 2006). There are two widely known resistance-conferring haplotype variants of *Rhg1* in soybean. The haplotype *rhg1-a* is commonly derived from “Peking” while *rhg1-b* is often derived from “PI 88788”, with the latter being used in over 95% of all commercial nematode resistant varieties (McCarville et al. 2017; Niblack et al. 2006). The Peking-type *rhg1-a* haplotype is less effective if it is not coupled with resistance-associated alleles at the unlinked *Rhg4* or *rhg2* loci (Basnet et al. 2022; Brucker et al. 2005; Yu et al. 2016). SCN populations that at least partially overcome three reference sources of *rhg1-b* resistance are designated as HG type 2.5.7. Throughout soybean-producing areas of the U.S., HG type 2.5.7 and other *rhg1-b*-virulent SCN populations are increasingly common and are gradually increasing their virulence on *rhg1-b* (Kleczewski et al. 2023; McCarville et al. 2017). In response, expanded use of Peking-type resistance and introduction of varieties that carry PI 88788-type *rhg1-b* resistance stacked with additional minor-effect resistance QTLs is anticipated (Bent 2022; Brzostowski and Diers 2017; Tylka et al. 2012)

The *rhg1-a* and *rhg1-b* haplotypes both carry three tightly linked genes that contribute to SCN resistance (Cook et al. 2014; Cook et al. 2012a). *Glyma.18g022400*, *Glyma.18g022500* and *Glyma.18g022700* respectively encode AAT_Rhg1_ (putative amino acid transporter), α-SNAP_Rhg1_ (vesicle trafficking mediator, also called GmSNAP18) and WI12_Rhg1_ (unknown function, carries a WI12 wound inducible protein domain). While the causal *Rhg1* genes were identified a decade ago and their mechanism(s) of action are being investigated, the resistance has been deployed for multiple decades. The α-SNAP_Rhg1_ proteins differ between the *rhg1-a* and *rhg1-b* haplotypes, as does the copy number of tandem repeats of the entire locus (Cook et al. 2014). The abundance of resistance-associated α-SNAP_Rhg1_ protein is known to increase 10-15-fold in the syncytium, the SCN feeding site (Bayless et al. 2016; Bayless et al. 2019). AAT_Rhg1_ protein increases an order of magnitude along the SCN root penetration path, where a substantial pool of AAT_Rhg1_ is associated with macrovesicles (Han et al. 2023). Less is known about how the WI12_Rhg1_ interacts with SCN, although impacts on root gibberellin levels have been discovered (Dong and Hudson 2022). Silencing or knocking out any one of the three genes increases susceptibility to SCN (Cook et al. 2012b; Dong and Hudson 2022; Liu et al. 2017; Schumacher and Bird 2025). In transgenic potato, expression of the set of three soybean *Rhg1* genes from their native soybean promoters or from strong constitutive promoters elevated resistance to potato cyst nematodes (Butler et al. 2019).

We hypothesized that *Rhg1* could also provide some resistance to Pp and that the commercially popular *rhg1-b* may have been beneficial against Pp. Melakeberhan and Dey previously reported that the infection rate of SCN in some instances decreased with increasing Pp proportions, and that the rate of Pp infection increased when co-inoculated at a 1:1 ratio with *H. glycines* but did not increase at higher or lower co-inoculation ratios (Melakeberhan and Dey 2003). That work was done using an SCN-susceptible soybean cultivar. Schumacher and Bird recently reported some instances where the presence of Pp increased SCN population densities on *rhg1-a* resistance sources, but with no overall consistent trend (Schumacher and Bird 2025). Interactions between *Rhg1* and other pathogens are only beginning to receive study, although most soybeans grown in recent years across the U.S. carry this genetic resistance (Lopez-Nicora et al. 2023). On soybean, Niblack and colleagues previously found that resistance to SCN was not reduced during co-infections with the root knot nematode *M. incognita* (Niblack et al. 1986). Significantly, the *rhg1-a* + *rhg2 (GmSNAP11)* combination has recently been shown to enhance quantitative resistance against reniform nematodes (Usovsky et al. 2021; Usovsky et al. 2022)

In this study we assessed (i) the success of Pp and an SCN HG 2.5.7 population on the soybean lines IL3025N (*rhg1-b*), IL3849N (*rhg1-a/Rhg4*) and Williams 82 (SCN-susceptible *rhg1-c*), in terms of total nematodes recovered per plant at the end of 30 days, (ii) the impact on nematode numbers when SCN and Pp were co-inoculated at 2:1 or 1:1 SCN:Pp ratios, and, using a set of near isogenic lines (NILs) for *rhg1-b*, (iii) how *rhg1-b* impacts Pp and SCN nematode numbers and the transcript abundance response of the three *rhg1-b* genes as well as salicylic acid (SA) and jasmonic acid (JA) response indicator genes.

## Materials and Methods

### Sand cup preparation

Quarried sand from Dane County, Wisconsin was sieved with a 0.25mm screen to remove larger gravel or sand particles and the resulting fine sand was moistened, placed in Nalgene polypropylene heat-tolerant autoclave bins at a maximum of 7.5 cm depth, and heated in a 65 °C oven for 48 h. The sand was cooled to room temperature, filled in plastic pots (9 cm deep, 6×6 cm at top tapering to 4×4 cm with drainage holes<u>, filled with 190 +/- 10 cc of sand</u>) and each of these sand cups (pots) was saturated with 50 mL of tap water.

### Plant genotypes

Seeds of Williams 82 (Wm82; *rhg1-c*), IL3025N (*rhg1-b*) and IL3849N (*rhg1-a* and *Rhg4*, susceptible allele at *Rhg2*) were used in the first set of cup assays. <u>Wm82 is a standard SCN-susceptible check; soybean lines that carry *rhg1-b* exhibit strong resistance to HG 0 SCN but only quantitative/partial resistance to HG 2.5.7 SCN; lines with *rhg1-a* + *Rhg4* exhibit strong resistance to HG 0 SCN and HG 2.5.7 SCN (Bent 2022; Niblack et al. 2006).</u> Seeds of Wm82 and near-isogenic lines Resistant NIL (Res. NIL; *rhg1-b*; LD09-15604) and Susceptible NIL (Sus. NIL; *rhg1-c*; LD09-15630) were used in the second set of cup assays and differential gene expression studies. IL3025N (synonyms: Illini 3025N, LD11-2170) and IL3849N (synonyms: Illini 3849N, LD07-3395bf) were developed by Brian Diers, University of Illinois at Urbana-Champaign, and are available through Baird Seed Co. (Williamsfield, Illinois). The pedigree for IL3025N is Syngenta_03JR313108 x LD05-3171. IL3849N is an SCN-resistant reselection from LD07-3395 whose pedigree is Syngenta_WW115926 x LD00-2817. The NILs were developed by Brian Diers, University of Illinois at Urbana-Champaign, through four backcrosses using LD00-3309 (from Maverick x Dwight, carries *rhg1-b*) as the recurrent parent and IA3023 as donor of the susceptible *rhg1-c* allele, hence the resulting NIL accessions are predicted to be >96% LD00-3309 genotype. All seeds were surface sterilized using chlorine vapor as described by Clough and Bent (1998).

### Nematode inoculum

The Pp population was collected in Wisconsin and maintained on ‘IO Chief’ sweet corn (*Zea mays*) root explants by inoculation onto 1-week old roots from freshly germinated seed grown on 1X Gamborg’s B-5 medium with 2% sucrose and 1.5% agar (Rebois and Huettel, 1986). Pp subcultures were started 3 months prior to use in experiments. The “SCN HG 2.5.7 IL” field population, obtained courtesy of Alison Colgrove, was maintained on IL3025N soybean plants and exhibited an FI ∼70 <u>on that *rhg1-b* soybean host</u> relative to reproduction on Williams 82.

SCN eggs from crushed cysts were hatched in 4 mM ZnCl_2_ solution at 27°C and dark conditions. Freshly hatched J2 stage SCN were rinsed and re-suspended in de-ionised water immediately prior to inoculation. Pp (all life stages other than eggs/J1) were collected together from corn root explants using the Baermann funnel method (Baermann 1917). The Pp were then rinsed and resuspended in de-ionised water. Pp inoculum batches had inherent variation but did consistently exhibit a demographic representation of, very approximately, 35% adult females, 15% adult males, 40% third and fourth stage juveniles and 10% second stage juveniles. Within each experiment the Pp aliquots applied to each of the Pp-inoculated plants were highly similar because all aliquots were drawn from the same Pp nematode preparation. Nematode inoculum density was adjusted so that 2 ml of liquid (1 ml containing Pp or water + 1 ml containing SCN or water) was inoculated into the sand of each pot via a small hole made using a 200 uL pipette tip, parallel to and 1 cm away from the soybean seedling, which was then re-covered with sand. The plants remained covered with humidity domes for one day post inoculation.

### Nematode count cup assays

<u>Unless otherwise noted, three nematode treatments were included in each experiment: SCN alone, Pp alone, or SCN + Pp.</u> For the experiments presented in Figure 1, four independent experiments were conducted, two using a 1:1 ratio of SCN:Pp (500 of each) and two using a 2:1 SCN:Pp ratio (approximately 1500 SCN:750 Pp) for co-inoculations. <u>Other pots in these experiments received only one nematode species (500 of either SCN or Pp in the 1:1 experiments; 1500 SCN or 750 Pp in the 2:1 experiments)</u>. <u>Each pot contained ∼190 cc of sand (described above).</u> The 2:1 experiments were conducted one week apart using separate SCN hatches but <u>used Pp from the same Baermann funnel extraction</u>. <u>The two 1:1 experiments each used independent preparations of SCN and Pp and were conducted {X} months apart from the eachother and separate from the 2:1 experiments.</u> Within each experiment there were five replicates of each treatment (3 plant genotypes x 3 nematode treatments = 9 treatments per block), arranged in a randomized complete block design<u>. The three plant genotypes (SCN-susceptible Wm82, SCN-resistant IL3029N and partially SCN-resistant IL3025N) are described above</u>. Each block of 9 pots was in a separate tray with a different randomized arrangement, with trays adjacent to each other in a large controlled environment growth room, and harvest\processing was done by block. Seeds were germinated in paper towels at 27°C for 4-5 days in the dark until a 5 cm radicle had formed and were then transplanted with one plant per sand pot. Nematode inoculum (see previous section) was applied one day after transplanting these VE stage seedlings (Vegetative Emergence) (Pedersen 2009). Plants were covered with a humidity dome for the first 2 days after transplanting. Each experiment was done twice (two separate dates). Inoculated plants were maintained in a growth room with 14h daylight, day temperature 27°C, night temperature 22°C, for 30 days. The plants were watered minimally (liquid did not flow from holes into the tray) by applying 10 mL of tap water to each pot each day for the first week, 15 mL a day the second week, alternating between 15 mL and 30 mL a day the third week and 30 mL a day thereafter. The pots were fertilized once after 15 days and again 7 days later using MiracleGro (Scotts, Marysville OH) as per manufacturer recommendations instead of tap water.

**Figure 1.**
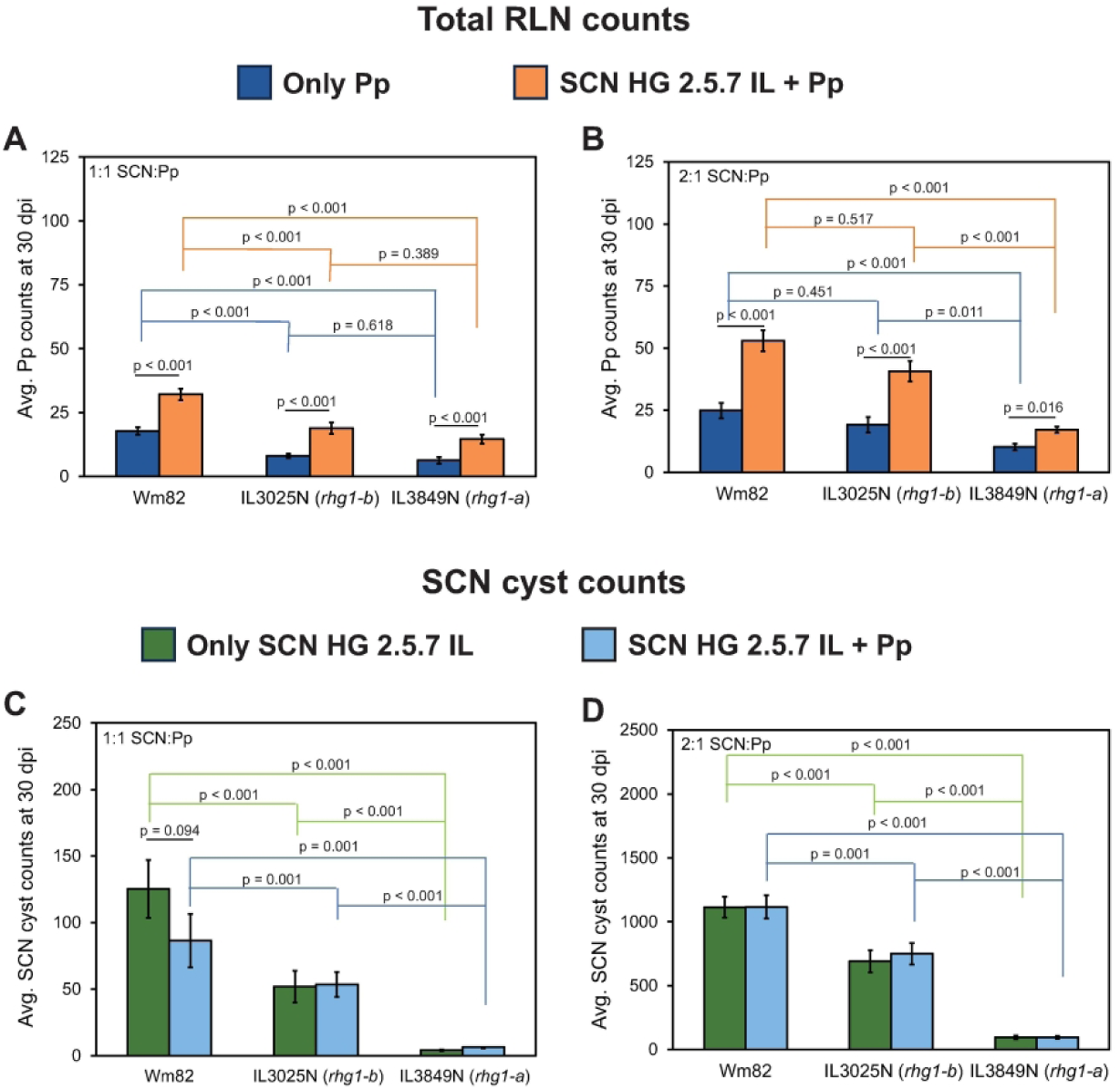
Nematode counts on Wm82, IL3849N (*rhg1*-a) and IL3025N (*rhg1*-b) plants at 30 dpi with either or both SCN HG 2.5.7 IL and Pp. (A) Pp counts at 30 dpi at 1:1 SCN: Pp inoculation (B) Pp counts at 30 dpi at 2:1 SCN:Pp inoculation. Blue bars correspond to Pp-only inoculation and orange bars represent combined inoculation of both SCN and Pp. (C) Numbers of SCN cysts recovered at 30 dpi at 1:1 SCN:Pp inoculation. (D) Numbers of SCN cysts recovered at 30 dpi at 2:1 SCN:Pp inoculation. Green bars represent SCN-only inoculation and the light-blue bars represent combined inoculation of both SCN and Pp. Values shown are mean +/- standard error of mean for raw (non-transformed) nematode counts. For the comparisons of greatest interest, statistical significance for differences between transformed means are displayed as p-values from ANOVA Tukey HSD tests (see Methods). The SCN HG 2.5.7 IL used partially overcomes *rhg1-b* resistance and represents the most prevalent HG type in the USA. Data are for two independent experiments with five replicates within each experiment (n = 10 for each bar).

For the experiments presented in Figures 2 and 3, a similar experimental design was used_-<u>(3 plant genotypes x 3 nematode treatments = 9 treatments per block, arranged in a randomized complete block design;</u> (two independent experiments with five replicates within each experiment, using distinct batches of seedlings and fresh nematode inoculum in each experiment<u>). The three plant genotypes in this case were standard SCN-susceptible check Wm82, SCN-susceptible Sus. NIL and partially SCN-resistant (*rhg1-b*) Res. NIL, treated with</u> approximately 500 <u>SCN, 500 Pp or 500 SCN + 500 Pp.</u>

**Figure 2.**
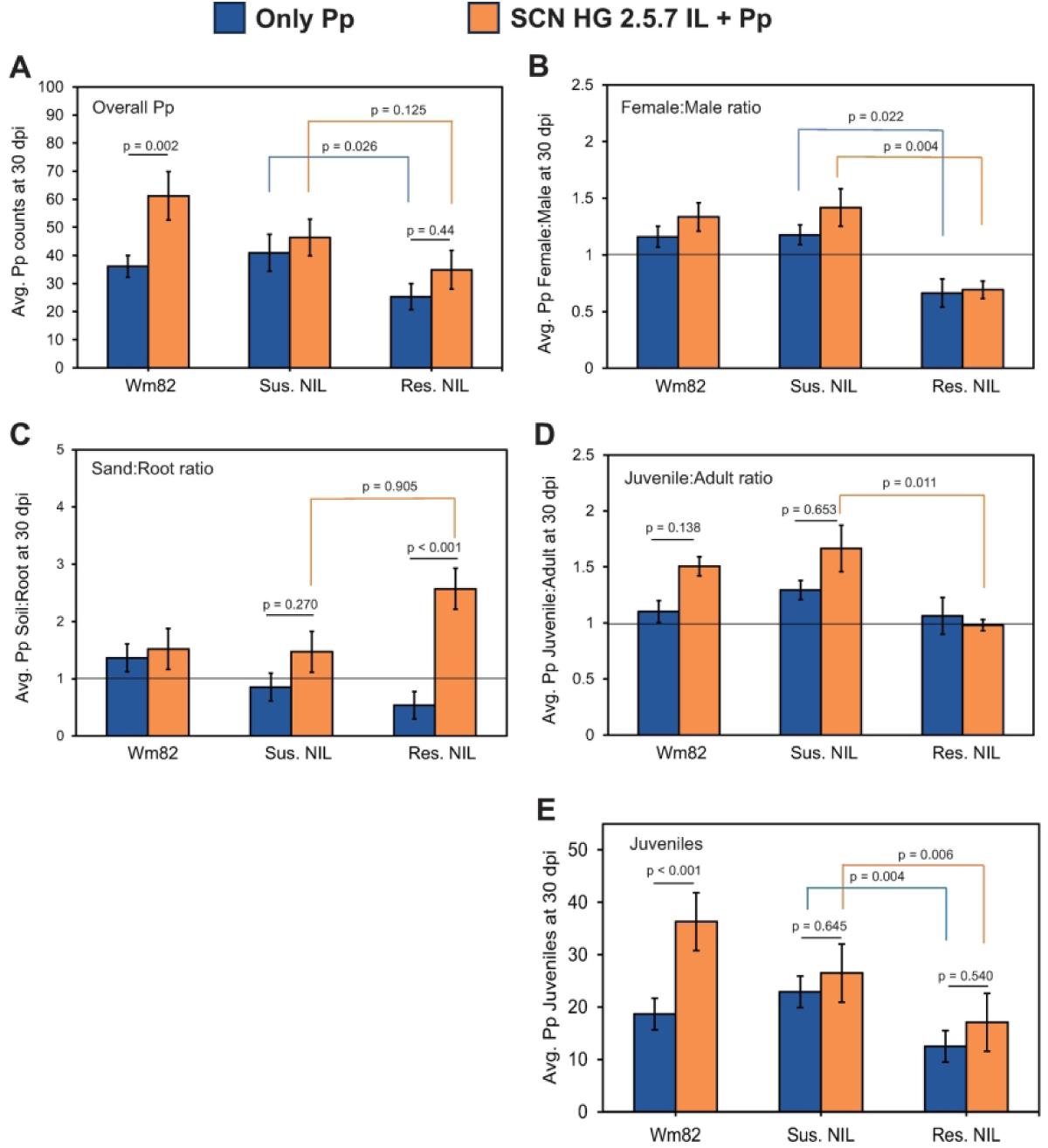
Pp counts on Wm82, Sus. NIL and Res. NIL plants at 30 dpi in either Pp-only inoculations (blue bars) or a combination of SCN and Pp at 1:1 ratio (orange bars). (A) Counts of all Pp. (B) Ratio of Pp females to males. (C) Ratio of Pp in sand fraction to root fraction. (D) Ratio of Pp juveniles (J1 through J4) to adults (male + female). (E) Counts of Pp juveniles (J1 through J4). Values shown are mean +/- standard error of mean for non-transformed data. Instances of possible significant differences of means are marked with p-values from ANOVA Tukey HSD tests performed on transformed data (see Methods). p > 0.2 if no p value is shown. The SCN HG 2.5.7 IL used partially overcomes *rhg1-b* resistance and represents the most prevalent HG type in the USA. Data are for two independent experiments with five replicates within each experiment (n = 10 for each bar).

**Figure 3.**
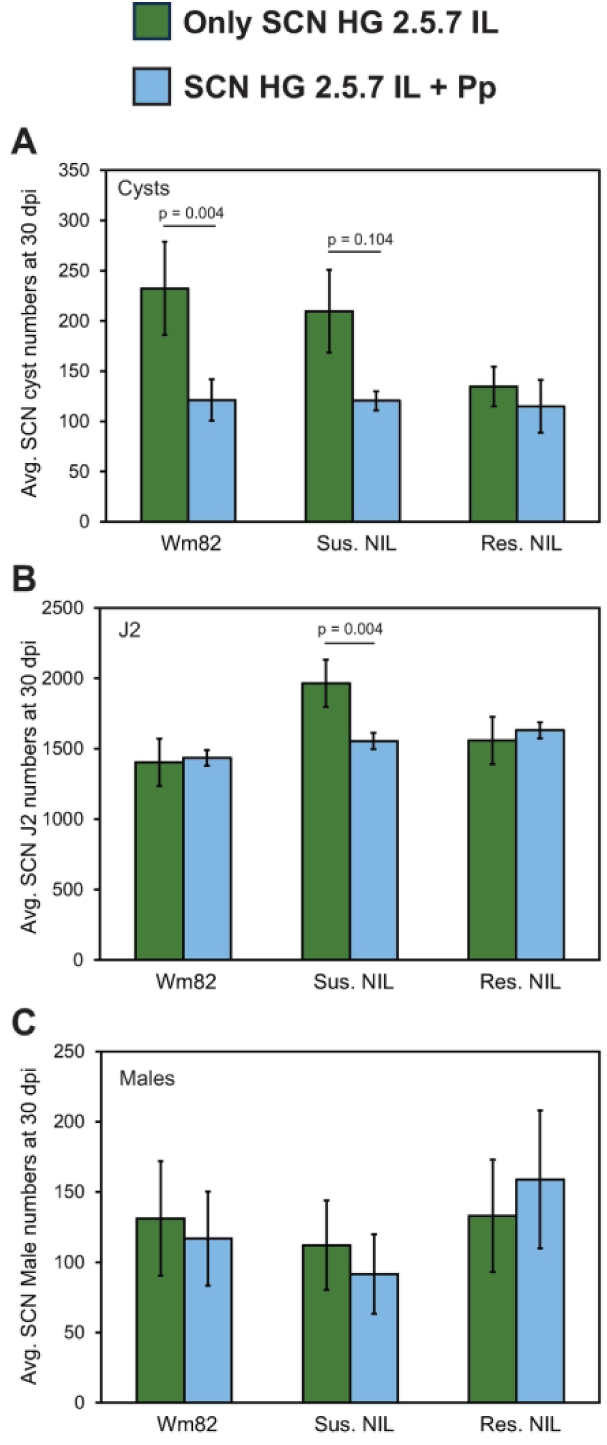
SCN cyst counts on Wm82, Sus. NIL and Res. NIL plants at 30 dpi in either SCN-only inoculation (green bars) or a combination of SCN and Pp at 1:1 ratio (light-blue bars). (A) SCN cyst counts. (B) SCN J2 counts (sand + root). (C) SCN male counts (sand + root). Values are mean +/- standard error of mean for non-transformed data. Instances of possible significant differences of means are marked with p-values from ANOVA Tukey HSD tests performed on transformed data (see Methods). p > 0.2 if no p value is shown. The SCN HG 2.5.7 IL used partially overcomes *rhg1-b* resistance and represents the most prevalent HG type in the USA. Data are for two independent experiments with five replicates within each experiment (n = 10 for each bar).

After 30 days, nematodes were extracted from the sand using water flotation and decanted into a nested sieve system consisting of a 2800 μm, 710 μm, 177 μm and 38 μm sieve (ASTM E11 US Alternative sieves 7, 25, 80, and 400 respectively). Plant roots and other material retained on the 2800 μm sieve after vigorous rinsing with tap water were collected, then the roots of all samples were chopped into ∼1 cm pieces and incubated on Baermann funnels with de-ionised water for 48 h at room temperature (25°C). 50 mL of water was eluted from the bottom of the funnels and evaluated for Pp (all life stages other than eggs/J1) and SCN J2s and males. Contents retained by the 177 μm sieve were <u>enumerated</u> for SCN cyst<u>s</u> under a stereoscope while contents from the 38 μm sieve were enumerated for Pp of all life stages (juveniles and adults) and for SCN males and juveniles. Nematode samples were further clarified by dilution using 70% (70 g into 100ml H_2_O) sucrose solution prior to microscopic enumeration. In pilot tests with no plants, average Pp extraction efficiency 24 h after inoculation was 54% (std.dev. 1.1%) using the above protocol. In NIL samples co-inoculated with both nematodes, contents from 38 μm sieve were also evaluated for SCN J2 and male numbers.

### NIL *rhg1* differential mRNA abundance assay

Each mRNA abundance result reports data from two independent experiments that used distinct batches of seedlings and nematode inoculum. Within each experiment that used roots from VE seedlings, three replicate blocks were set up in a randomized complete block design as described above for nematode count cup assays except with two pots of each treatment within each block, to allow sampling at the 3 dpi (days post inoculation)and 7 dpi time points. Note that infection can be asynchronous and these time points can include infections initiated 0-3 days or 0-7 days prior to sampling. For gene expression experiments that used V1 (Vegetative 1^st^ stage) plants (Pedersen, 2009), inoculation occurred when the first set of trifoliate leaves had emerged and two blocks were utilized. For RNA extraction, roots from plants (Wm82, Res. NIL and Sus. NIL) inoculated with either SCN (500 J2) or Pp (500 J2-to-adult worms) or both (500 SCN + 500 Pp) as well as non-inoculated controls were rinsed in water and observed under a stereoscope at 3 dpi or 7 dpi. Each sample was from a separate plant. For each sample, about 100 mg of root tissue<u>s</u> <u>from a single plant,</u> from ∼1 cm <u>segments</u> with<u> multiple</u> visible <u>ater-soaked foci (putative infection sites</u>, <u>were</u> placed in ice-cold cryovials with 5 ceramic beads and immediately frozen in liquid nitrogen. These samples were stored at -80°C. Infection sites were in the zone of maturation on lateral roots and similar root sites were collected from non-inoculated control root systems. Frozen tissue was processed at 2500 rpm for 30 s in a powerlyzer and returned to liquid nitrogen for a minute. This process was repeated 2-3 times until tissue was pulverized. RNA was then extracted from pulverized root tissue using Zymo (Irvine, CA) Direct-Zol TriZol-based miniprep kit using manufacturer protocol. cDNA was synthesized from 1µg of RNA for each sample using Solis BioDyne (Tartu, Estonia) FireScriptTM RT cDNA Synthesis mix with included Olido dT random degenerate hexamer primers (Catalog #06-20-00500), according to manufacturer protocol.

Real-time quantification and Ct values were obtained for transcripts encoding AAT_Rhg1_, WI12_Rhg1_, α-SNAP_Rhg1_HC (detects only the HC/ *rhg1-b* haplotype), as well as for *AAT* (*Glyma.10G109500*; JA response indicator), *NIMIN1* (*Glyma.10G010100*; SA response indicator) and the internal standard gene *NREG1* (*Glyma.11G079600*) (Beyer et al. 2021). ΔCt values were the average of two technical replicates from the same sample. To determine the fold-change in transcript abundance in response to nematode inoculations the 2^-ΔΔCt method was used (Livak and Schmittgen 2001). <u>For comparisons within plant genotypes,</u> ΔCt values obtained using NREG1 as internal standard were normalized to the average ΔCt values of all no-nematode samples for the same plant genotype within the same experiment date to obtain ΔΔCt values. <u>comparisons between</u>plant genotypes, the ΔCt values for all genotypes and treatments were normalized to the average ΔCt values of no-nematode inoculated Sus. NIL plants to obtain ΔΔCt values. A full list of the oligonucleotide primers used in this study is available in Supplementary Table 1.

### Statistical analyses

All data were analyzed and visualized using R (version 4.3.0). Graphs show mean and standard error of mean for the original non-transformed data. For statistical tests, nematode count data were square root transformed or log transformed to achieve normality (residuals checked graphically to confirm equal variance and normality). ANOVA was performed using the aov() function and the means were separated using TukeyHSD() in the package stats(). Plant genotype, nematode treatment, experiment, and blocks within experiment were all fixed effects and the interaction between genotype and nematode treatment was included in the model. The specific R model used is provided for each analysis in the Supplemental Tables. A typical analysis assessed one nematode type (e.g., Pp numbers in Pp-only and Pp+SCN treatments) and had 60 entries (2 experiments, 5 replicate blocks per experiment, 3 plant genotypes, 2 treatments). For the transcript abundance evaluations, experiments were set up similarly to the above and the same model was fit as for nematode counts (models stated in Supplementary Tables). Transcript statistical analyses were performed on the ΔΔCt values (mRNA abundance).

## Results

### Pp parasitism of soybean is enhanced by co-infection with SCN

Controlled environment single-species inoculation and co-inoculation experiments with Pp and SCN were carried out (Pp-only, SCN-only or Pp+SCN all within the same experiment). Experiments were first conducted using soybean genotypes that differ at hundreds of loci, including different SCN resistance haplotypes. Wm82 (*rhg1-c*) is SCN-susceptible while IL3025N (also known as LD11-2170) carries *rhg1-b* and IL3849N (also known as LD07-3395bf) carries *rhg1-a* and *Rhg4*. Four independent experiments were conducted, two using a 1:1 ratio and two using a 2:1 SCN:Pp ratio for co-inoculations, with five replicates of each treatment within each experiment. The experiments utilized an SCN HG 2.5.7 population that, as expected, reproduced well on the susceptible *rhg1-c* (Wm82) and substantially overcame *rhg1-b* resistance (IL3025N) while exhibiting poor reproduction on a plant line carrying the *rhg1-a* and *Rhg4* combination (IL3849N) (green bars in Figure 1C and 1D; all differences between SCN-only inoculations (green bars) were significant at p<0.001). ANOVA tables associated with this and subsequent figures are provided as Supplementary Tables.

Figure 1A and 1B show that more Pp were recovered upon co-inoculation with virulent SCN HG 2.5.7 IL compared to the Pp-only inoculations, in experiments on SCN-susceptible Wm82 or moderately SCN-susceptible IL3025N soybean plants (orange vs. blue bars for same soybean genotype, p < 0.001). More Pp were also recovered upon co-inoculation with SCN on IL3849N (Figure 1A and 1B), which is a genotype that largely prevents SCN cyst production (Figure 1C and 1D). The above observations were true for SCN:Pp inoculation ratios of 2:1 and 1:1. In all of the experiments in Figure 1, about twice as many Pp were recovered 30 days after inoculation when Pp were co-inoculated with SCN HG 2.5.7, compared to Pp-only inoculations (the ratio on the same plant genotype in the same experiment ranged from 1.7 to 2.3, across all plant genotypes and both ratios of co-inoculation).

Within the same experiments as above, there were no significant differences in SCN HG 2.5.7 IL cyst numbers between SCN-only inoculations and co-inoculations with Pp on the same plant genotype (green vs. blue bars, Figure 1C and 1D). Nearly 10-fold more cysts were recovered across treatments when ∼1500 (Figure 1D) rather than 500 (Figure 1C) SCN J2s were inoculated, for both solo inoculations and co-inoculations with Pp.

### *rhg1-b* impedes the parasitism of Pp

A separate phenomenon was observed in the experiments of Figure 1, suggesting that soybean SCN resistance might also impact RLN. When inoculated by itself or together with SCN, significantly fewer Pp were recovered after 30 days from the SCN-resistant variety IL3849N than from Wm82 (compare bars of same color in Figure 1A and 1B). Pp abundance on IL3025N was lower than on Wm82 in three of the four comparisons (Figure 1A and 1B). Fewer Pp were recovered from IL3849N than from IL3025N only for the 2:1 SCN:Pp inoculations (Figure 1B vs. 1A). However, the genetic backgrounds of these soybean genotypes differ at thousands of loci.

A more rigorous test for an effect of the SCN resistance *rhg1-b* haplotype on Pp was conducted by using near-isogenic soybean lines (NIL) expected to carry the same genotype at >96% of all loci but carrying either *rhg1-c* (Sus. NIL) or *rhg1-b* (Res. NIL) at the *Rhg1* locus. The results supported the conclusion that *rhg1-b* can negatively impact Pp (Figure 2). Significantly fewer Pp were recovered from the Res. NIL compared to the Sus. NIL when singly inoculated with Pp (Figure 2A blue bars). When co-inoculated with both nematodes (Figure 2A orange bars), fewer Pp were again recovered from Res. NIL plants than Sus. NIL plants but with a difference that was at the margin of statistical significance (p = 0.056).

For SCN, multiple studies have shown that the proportion of female to male nematodes formed is often lower on soybean plants with effective SCN resistance than on susceptible plants (Colgrove and Niblack 2005). Intriguingly, with migratory Pp we also observed that the ratios of females to males recovered from the *rhg1-b* Res. NIL were significantly lower than those for *rhg1-c* Sus. NIL plants (Figure 2B). This was true in both Pp-only and SCN co-inoculated treatments. When the proportion of Pp found in roots vs. the surrounding sand at 30 dpi was enumerated, the *rhg1-b* Res. NIL had a higher proportion of the Pp in the sand than the Sus. NIL, but intriguingly, that was only in the Pp + SCN co-inoculated samples (Figure 2C). For the Res. NIL line, a ∼5-fold higher proportion of Pp were in the sand than in the root upon co-inoculation with SCN, compared to the Pp-only samples (Figure 2C). The relative presence of juvenile Pp 30 days after inoculation was examined as another parasitism/resistance metric, because those juveniles are predominantly progeny produced from the initial inoculum. At 30 days after inoculation the number of juvenile-stage Pp (Figure 2D) and the proportion of juvenile-stage vs. adult Pp (Figure 2E) again showed that presence of SCN shifted Pp population demographics. Juvenile Pp were significantly less common on *rhg1-b* Res. NIL plants compared to the *rhg1-c* Sus. NIL. For the juvenile:adult ratio they were less common on Res. NIL plants only when Pp and SCN were co-inoculated (Figure 2E). The findings of Figure 2 together indicate that *rhg1-b* can exert a negative impact on Pp, in the presence of SCN but also in the absence of SCN.

### Pp co-inoculation reduces SCN cyst numbers only in the susceptible *rhg1-c* lines

As expected, in SCN-only inoculations fewer SCN HG 2.5.7 IL cysts were recovered from the resistant *rhg1-b* NIL plants compared to the susceptible *rhg1-c* NIL plants or Wm82 (Figure 3A; data are from same experiments as Figure 2). Co-inoculation of Pp with SCN reduced the number of cysts recovered on the SCN-susceptible *rhg1-c* Wm82 and Sus. NIL lines (Figure 3A). The difference was statistically significant only for Wm82 (Figure 3A), but a similar trend was also observed on Wm82 in the experiments of Figure 1A (p = 0.094) and on Sus. NIL in the experiments of Figure 3A (p = 0.070). When the plants had partial resistance to SCN that reduced cyst production (Res. NIL), co-inoculation with Pp did not further reduce cyst numbers (Figure 3A). In the same experiments, the J2 and male SCN populations in roots and sand at 30 dpi were also enumerated. The results were not consistent across the two SCN-susceptible plant genotypes, as fewer total SCN J2s were recovered after co-inoculation with Pp only in the case of the Sus. NIL plants (Figure 3B). Co-inoculation with Pp did not reduce J2 SCN numbers for Wm82 and Res. NIL plants (Figure 3B). The number of SCN males recovered exhibited substantial variation, and no statistically significant differences were observed between plant genotypes or between plants inoculated with only SCN and those inoculated with both nematodes (Figure 3C).

### No significant changes in *rhg1-b* transcript abundance detected in response to nematode infection

In light of the observed interactions between SCN, Pp and *rhg1-b*, we investigated nematode induction of increases in transcript abundance of the three *rhg1-b* genes as well as indicator genes of SA-response and JA-response activity in soybeans, at 3dpi and 7dpi (Figures 4 and 5). As has previously been observed (Cook et al. 2014), *Rhg1* transcripts were approximately 5- to 20-fold more abundant for the high-copy *rhg1-b* genotype than for the single-copy *rhg1-c* genotype (Supplemental Figure 1). Transcript abundances are reported in Figures 4 and 5 as fold-change relative to the mean of the abundance for the non-inoculated control of the same plant genotype within the same experiment (ΔΔCt). Figure 4 A-F show that even with two independent experiments that each sampled infected zones from three independent root systems, we did not detect any significant nematode-induced transcript abundance of any of the *rhg1-b* genes when plants were inoculated at VE stage (emerging seedlings, cotyledons not yet separated). We also observed no significant differences with plants inoculated at the V1 stage (fully expanded trifoliate; two independent roots in each of two experiments; Supplemental Figure 2).

**Figure 4.**
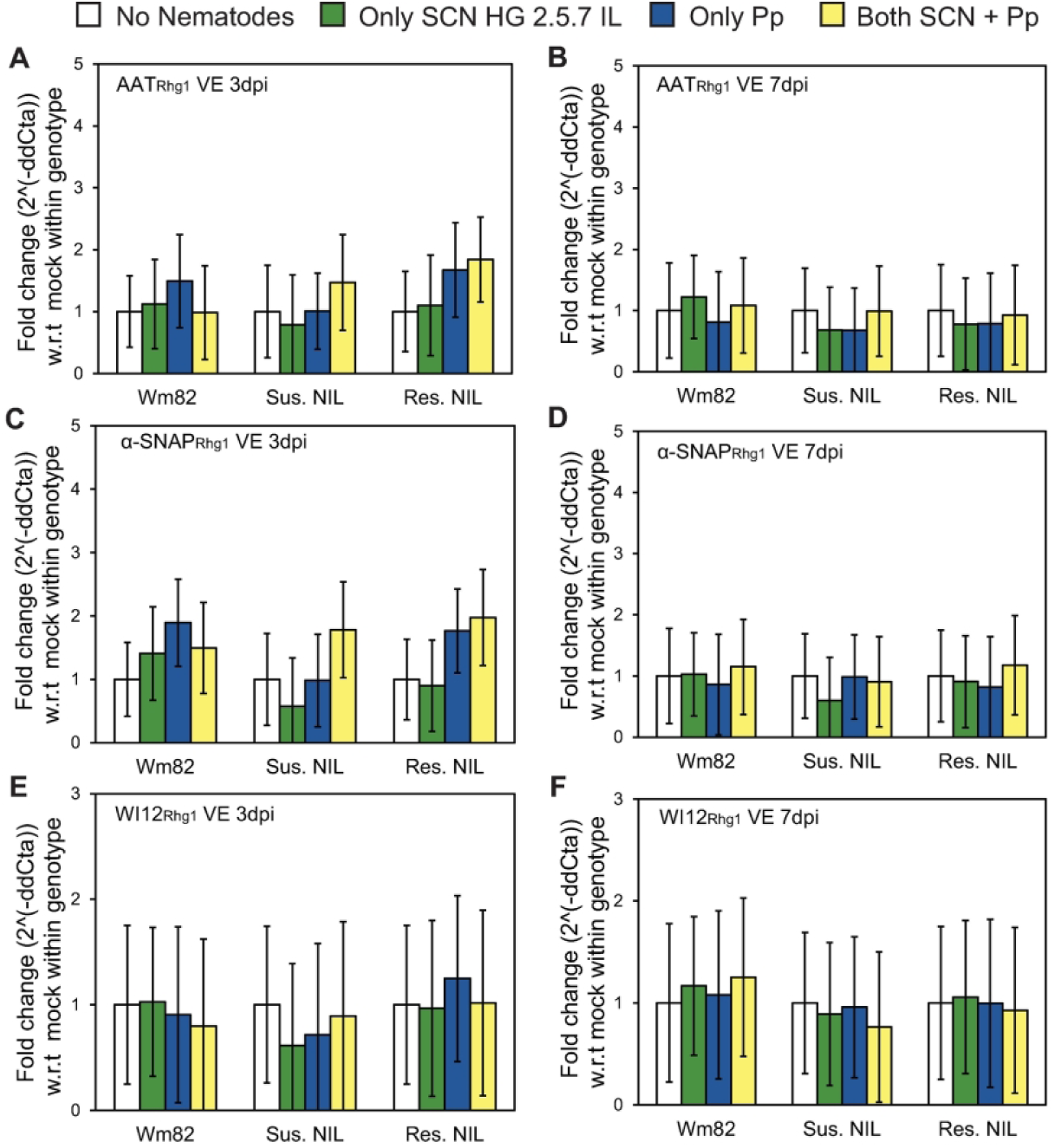
*Rhg1* transcript induction (fold change) relative to (w.r.t.) no-nematode controls within each plant genotype. The protein encoded by the analyzed transcript is noted in the upper-left corner of each graph. Wm82, Sus. NIL and Res. NIL plants were inoculated at VE stage and root samples from infection areas taken at: (A, C and E) 3 dpi, or (B, D and F) 7 dpi. No statistically significant differences between samples were observed using ANOVA and Tukey HSD test. The SCN HG 2.5.7 IL used partially overcomes *rhg1-b* resistance and represents the most prevalent HG type in the USA. Data are from two independent experiments with three replicate roots assessed within each experiment (n = 6 for each bar).

**Figure 5.**
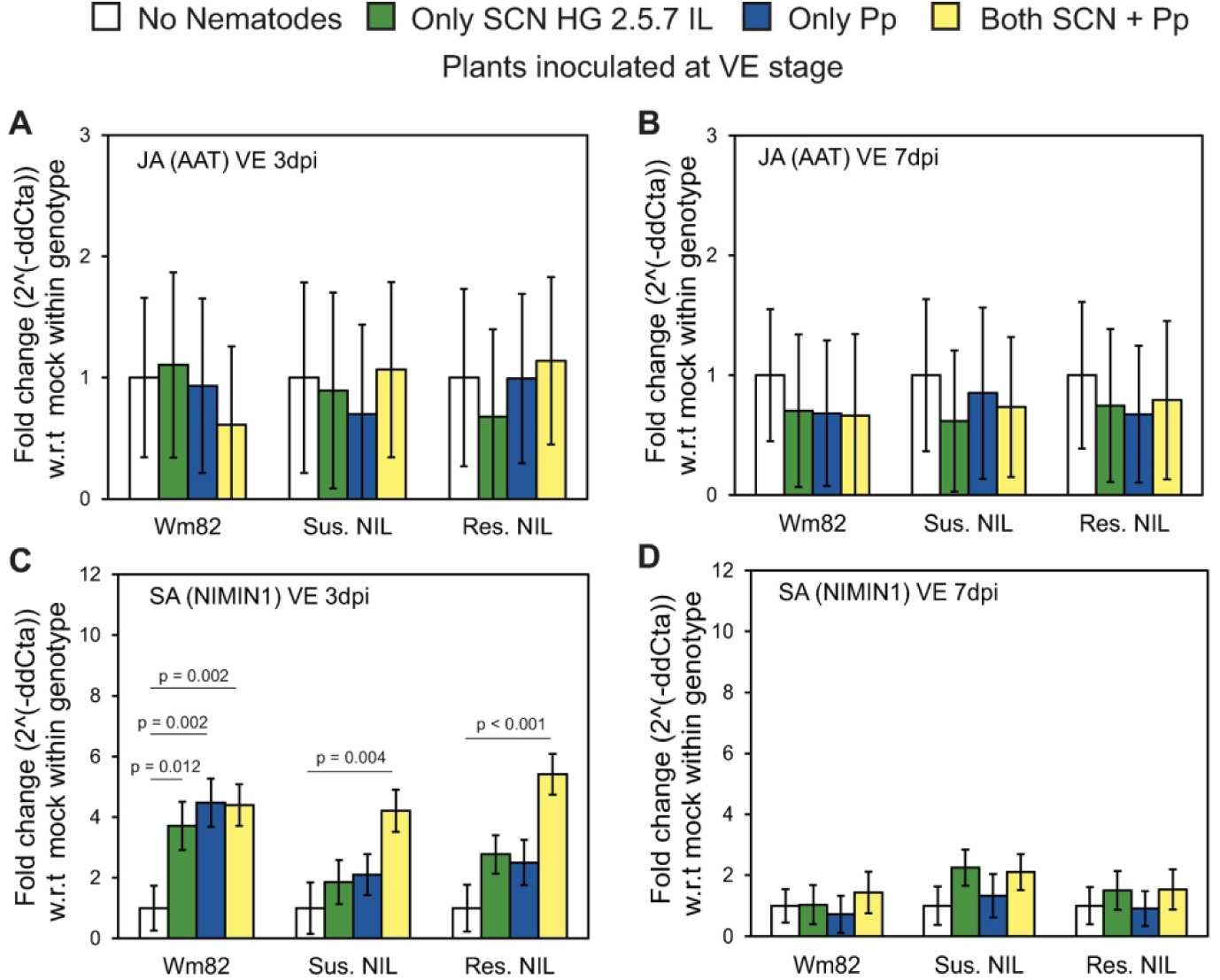
Induction (fold change) of JA-response indicator gene *Glyma.10G109500* and SA-response indicator gene *Glyma.10G010100* transcripts at 3 dpi and 7dpi, relative to (w.r.t.) no-nematode controls within each plant genotype within the same experiment. Wm82, Sus. NIL and Res. NIL soybean accessions inoculated at VE stage; transcript measured and sample dpi indicated in upper left corner of graph. Significant differences of means are marked with p-values from ANOVA Tukey HSD tests (p > 0.2 if no p value is shown). The SCN HG 2.5.7 IL used partially overcomes *rhg1-b* resistance and represents the most prevalent HG type in the USA. Data are from two independent experiments with three replicate roots assessed within each experiment (n = 6 for each bar).

### Combined infection by SCN and Pp induces a stronger salicylic acid response

At either 3 dpi or 7 dpi, neither SCN, Pp nor their combination induced a significant change in transcript abundance (ΔΔCt) for the JA-response indicator gene ‘*AAT*’ (Figure 5A and 5B). Note that this established soybean JA-response indicator gene *AAT* (Beyer et al. 2021) is *Glyma.10G109500* and is not the *Rhg1* gene for AAT_Rhg1_. In contrast to the JA result, transcript abundances for the soybean SA-response indicator gene *NIMIN1* (*Glyma.10G010100*) did exhibit statistically significant increases at 3 dpi in many of the nematode-inoculated samples (Figure 5C). Only dual inoculation with SCN and Pp consistently caused significant elevation of the SA indicator in all three soybean genotypes (Figure 5C). The elevated SA response indicated by *NIMIN1* transcript levels was no longer observable at 7 dpi (Figure 5D).

## Discussion

Although SCN is the most economically important nematode of soybeans in the US, in more tropical soybean producing states such as Brazil, RLN (especially *Pratylenchus brachyurus*) has been reported to be more consequential (Castanheira et al. 2020; Nomura et al. 2024). No soybean varieties labeled as having strong resistance to RLN have been deployed but searches are underway (e.g., Chowdhury et al. 2022; Hamawaki et al. 2019; Machado and Araúujo 2016). SCN and RLN frequently co-occur. Given the importance of the soybean crop, it is crucial to understand if there are interactions between the two nematodes on soybean and if that could lead to a change in their relative abundance over time. Only limited surveys and studies are available on the tripartite interaction of Soybean-SCN-RLN (Melakeberhan and Dey 2003). At the time of writing this manuscript, no published reports are available on how the deployed *Rhg1* resistance against SCN factors into this tripartite interaction.

Many soybean fields are infested with both SCN and RLN and the limited research reported suggests that their interaction is complicated. MacGuidwin et al. suggested a competitive relationship (population densities of both SCN and RLN suppressed by co-occurrence) based on an analysis of field survey data (MacGuidwin et al. 2023). Melakeberhan and Dey tested four 1000-nematode inocula with varying proportions of SCN and Pp and found evidence of competition when SCN outnumbered Pp 3:1. When both species were equally represented the interaction appeared to be antagonistic, as numbers of Pp increased and SCN numbers decreased as compared to the single-species case (Melakeberhan and Dey 2003). Those experiments used HG 0 SCN on an SCN-susceptible soybean host. Taking a cue from that prior study (Melakeberhan and Dey 2003), we co-inoculated equal numbers of SCN and Pp, and then also examined how twice as many SCN as Pp altered the result. <u>As in the study of Melakeberhan and Dey w</u>e observed that presence of SCN increased Pp numbers on soybean roots, but also observed the increase on an SCN-resistant host (IL3849N). In contrast to the prior study (Melakeberhan and Dey 2003), we observed more not fewer Pp 30 days after co-inoculation with twice as many SCN. Hypotheses for how SCN assists Pp parasitism include effector-mediated suppression host defenses, and/or simply by providing established entry points into the root.

We used HG 2.5.7 SCN both because it is informative with respect to *Rhg1* (it largely overcomes *rhg1-b* but not *rhg1-a* + *Rhg4*), and because it is becoming the most common type of SCN found in North Central U.S. soybean fields. The capacity of SCN to increase Pp numbers on IL3849N is a related topic for future study - Pp may for example still be quantitatively impacted by the reduced number of SCN present on the resistant host after the first few days of infection, and/or by the plant defense responses induced by SCN on the resistant host.

The Pp life cycle can require longer than 30 days. However, we were testing Pp together with SCN and hence performed 30-day experiments. 30 days is the established assay time for SCN resistance assays because longer time points introduce the issue of multiple generations of SCN reproduction. A priori, impacts on any plant-parasitic nematode can be documented without taking the study through a full nematode life cycle. However, future studies could incorporate a full life cycle assessment of Pp during co-infection with SCN.

The overall infection efficiency of Pp on soybean as opposed to other hosts also merits future study. Pp did infect soybean in the present study setting, and infected SCN-susceptible soybean lines more when there were more SCN present (Figure 1). At the 30-day time point it was often the case (see Figure 2C) that in the absence of SCN over half of the Pp were in the soybean roots and not in the sand media. We also were able to readily find water-soaked infection sites on the roots for our RNA extraction and qPCR experiments. Some attrition (nematodes that do not find and feed on the root and hence do not survive) is common in any nematode inoculation assay and attrition will increase as inoculum size per root increases. However, out of an initial inoculum of 500, the number of Pp recovered from roots and sand at 30 dpi was often below 10%. We speculate that, despite the care taken to water pots consistently and with small volumes, sub-optimal watering (that washes nematodes from the root zone and/or imposes low-moisture stress on the nematodes) may have been responsible for this low Pp recovery rate.

In our previous studies with *Rhg1* genes we had shown their capacity to increase resistance to other cyst nematode species and genera (*Heterodera schactii*, *Globodera pallida*, *Globodera rostochiensis*) when transgenically expressed in *Arabidopsis* or potato (Butler et al. 2019). Similarly, the combined presence of SCN resistance loci *rhg1-a* and *rhg2* was shown to also provide quantitative resistance against reniform nematodes (*Rotylenchulus reniformis*) (Usovsky et al. 2021; Usovsky et al. 2022). The present study was designed as an initial investigation of the broader hypothesis that SCN-resistant soybeans deployed in fields may have been partially controlling RLN. Our data are not inconsistent with the hypothesis that early (30 dpi) establishment of Pp can be impeded by *rhg1-a* + *Rhg4*, as evidenced by our observed reduction of Pp population sizes in IL3849N relative to Wm82. Early Pp establishment was also impeded on IL3025N (*rhg1-b*) when initial SCN and Pp population densities were similar, but was not significant when SCN outnumbered Pp. We further hypothesized that the interaction between SCN and Pp could reduce the efficacy of *Rhg1*. In support of that, as noted above, relative to inoculation of the same plants with just Pp we found an increase in Pp recovery when simultaneous SCN pressure was applied. Interestingly, for the virulent SCN HG 2.5.7 IL population that we tested, counts of SCN cysts were not reduced by co-inoculation with Pp except at 1:1 ratio when *rhg1-a* or *rhg1-b* were absent. A similar dynamic between SCN and Pp was observed by Melakeberhan and Dey on multiple soybean cultivars that lacked the *rhg1-a* or *rhg1-b* resistance (Melakeberhan and Dey 2003). We have not investigated full-season resistance or yield protection (see also Pedro et al. 2024), but the reductions in parasitism observed in the present study and that of Melakeberhan and Dey suggest a potential contribution to full-season disease resistance against Pp. The field studies of Saikai and MacGuidwin (Saikai and MacGuidwin 2022) were done using varieties with *rhg1-b* resistance, so a yield loss impact of Pp on SCN-resistant soybean varieties has been demonstrated. The present study suggests that Pp damage could be even worse if use of *rhg1-b* soybeans was not prevalent, but further field-level and full-season or multi-season studies are needed to investigate this point.

As noted above, in our experiments and those of Melakeberhan and Dey (2003) co-inoculation with SCN helped Pp or was neutral, in all cases tested. Even on resistant hosts, we observed no reduction of Pp at 30 dpi when co-inoculated with SCN. In contrast, with wheat, Laserre et al. 1994 and Mokrini et al. 2018 both observed that on susceptible hosts co-inoculation with cereal cyst nematode tended to reduce *Pratylenchus* population sizes (Lasserre et al. 1994; Mokrini et al. 2018). Intriguingly Laserre and colleagues also showed, on a susceptible host, a reduction in eggs and vermiform *P. neglectus* upon inoculation of *H. avenae* onto a separate part of the root system. However, on wheat accessions with resistance to *H. avenae* they observed that co-inoculation with *H. avenae* led in some cases to higher numbers of *Pratylenchus*, similar to the trend in our experiments (Lasserre et al. 1994).

The *rhg1-b* haplotype SCN resistance, unlike the *rhg1-a* haplotype resistance, is not co-dependent on the *Rhg4* locus on chromosome 08 (Patil et al. 2019). We were able to obtain near-isogenic lines for *rhg1-b* that, after four backcrosses, should be genetically >96% identical, for a more well-controlled test of the contribution of *rhg1-b* to resistance against Pp. As with IL3025N and IL3849N, we observed fewer Pp on Res. NIL plants compared to the Sus. NIL plants when only Pp was inoculated, strongly linking *rhg1-b* SCN resistance to partial Pp resistance as well. In addition to the overall reduction in Pp numbers in the Res. NIL, we also observed reductions in the ratios of females to males and juveniles to adults. RLN males and females differ in how they feed on the roots and how much damage they can subsequently cause (Saikai and MacGuidwin 2020). Females make larger lesions and burrow more frequently into the roots than the males. Hence having overall fewer Pp and a population that is skewed towards adults, especially males, may be less damaging to the crop. When considered together, these observations indicate that *rhg1-b* imposes stress on Pp populations and further support the indication that *rhg1-b* may be reducing the severity of damage to soybean caused by Pp.

In an earlier pair of studies, we had observed AAT_Rhg1_ protein abundance increasing along SCN penetration path (Han et al. 2023), unlike the α-SNAP_Rhg1_ that exhibited a strong abundance increase in the syncytium (Bayless et al. 2016). Because Pp do not form syncytia, in the present study we hypothesized that *Rhg1-GmAAT* could be one of the early response genes that functions against diverse nematodes. However, for the genes encoding AAT_Rhg1_, α-SNAP_Rhg1_ or WI12_Rhg1_ we did not observe any statistically significant transcript induction at 3 dpi for plants inoculated with Pp at the VE or the V1 stage, or at 7 dpi after VE stage inoculation. It is possible that any transcript induction is sufficiently local to infected cells that the vast majority of non-infected cells in the root samples diluted out any signal from cells at or immediately adjacent to the site of nematode infection. It is also possible that expression of the *Rhg1* gene products is regulated in part at the post-transcriptional level. This absence of an observed transcript induction is consistent with a previous study where we saw no significant differences in transcript induction for α -SNAP_Rhg1_HC at 7 dpi but did observe significant changes in protein abundance (Haarith et al. 2025).

Transcript abundances for *NIMIN1*, a preferred soybean salicylic acid response pathway indicator (Beyer et al. 2021), were upregulated significantly at 3 dpi in response to SCN, and were more reliably upregulated when Pp co-infected alongside SCN. Perhaps of greater interest, Pp alone did not reproducibly elicit this SA response while Pp+SCN did (Figure 5), and this Pp co-infection with SCN enhanced both SA responses and Pp population sizes (Figure 1). Antagonism between SA- and JA-mediated defense responses has been widely reported (Glazebrook et al. 2003; Verma et al. 2025). There are multiple instances of a “host mis-direction” virulence strategy in which pathogens or pests that are more effectively defended against by the one set of defense responses cause the plant to instead activate the other defense responses (Schweiger et al. 2014).

The non-sedentary (migratory) parasitism exhibited by Pp may be more similar to plant attack by herbivorous insects than it is to a haustorium-forming biotrophic fungus or oomycete, or a syncytium-forming cyst nematode. Our results suggest that Pp may lose more to JA-mediated plant defenses than to SA-mediated plant defenses; future studies could investigate if Pp in fact benefits from stronger elicitation of SA-mediated defenses.

The findings of (Usovsky et al. 2021) correlating α-SNAP-encoding genetic loci with resistance to both SCN and reniform nematode support speculation that α-SNAP protein variants may also contribute to resistance against Pp. A large body of evidence has implicated SNAP proteins in SCN resistance, including major roles for the α-SNAP_Rhg1_HC/GmSNAP18 protein encoded by *rhg1-b* and the α-SNAP_Rhg1_LC/GmSNAP18 protein encoded by *rhg1-a*, and more subtle contributions from a γ-SNAP (gamma-SNAP) encoded at *cqSCN-006* (*Glyma.15G191200*) and loss of the α-SNAPs encoded at *GmSNAP02* (*Glyma.02G260400*), and *rhg2*/*GmSNAP11* (*Glyma.11g234500*) (Bayless et al. 2016; Bent 2022; Butler et al. 2021; Cook et al. 2014; Cook et al. 2012a; Haarith et al. 2025; Lakhssassi et al. 2017; Liu et al. 2017; Matsye et al. 2012; Usovsky et al. 2023). Four separate α-SNAP proteins bind to a single SNARE bundle as they mediate NSF binding and SNARE bundle disassembly (Zhao et al. 2015), and presence of multiple types of SNAP proteins is likely to impact their function. Future research could reveal that certain combinations of SCN resistance-associated α-SNAP-encoding loci may provide stronger Pp resistance than *rhg1-b*, and/or provide diversified resistance mediators that impose distinct selection pressures on Pp populations. However, a leading hypothesis for the impact on cyst nematodes of the non-canonical SCN resistance-associated α-SNAPs is their cytotoxicity and elevated abundance in syncytia (Bayless et al. 2016; Bayless et al. 2018), and Pp are not sedentary feeders that induce syncytia. Some Pp individuals do feed on a single host cell for several hours (Zunke 1990), which may be enough time for SCN resistance-associated SNAPs to perturb the susceptible status of the host cell. But the *Rhg1*-encoded AAT_Rhg1_ and WI12_Rhg1_ proteins also contribute to SCN resistance (Cook et al. 2012). An equally strong hypothesis for future research is that AAT_Rhg1_ and/or WI12_Rhg1_, moreso than α-SNAP_Rhg1_, are the primary mediators of the impeded Pp parasitism caused by *Rhg1*.

Taken together, the present results indicate that the *rhg1-b* haplotype impedes the parasitism of Pp as well as SCN – a pair of plant parasitic nematode species that are highly distinct taxonomically and behaviorally. The present controlled environment results also show, in agreement with a past study, that on SCN-susceptible soybean genotypes SCN can enhance Pp population sizes. There is measurable interaction between the two nematodes regarding the efficacy of *rhg1-b* resistance, and in their induction of salicylic acid responses within the plant. The findings suggest multiple avenues for future study and manipulation of soybean resistance to *Pratylenchus* root lesion nematodes.

## Supporting information

All supplemental information

## Author Contributions

DH conceptualized, designed, and performed the experiments, analyzed the data, and authored the manuscript. AEM provided Pp inoculum, methods and expertise in *Pratylenchus* biology and edited the manuscript. AFB co-designed the experiments, and analyzed and co-authored the manuscript. AFB and DH are co-PIs on the grant that funded this study.

## Acknowledgements

Funding for this study was provided primarily by a grant to AFB and DH from the Wisconsin Soybean Marketing Board, and also by grants to AFB from the United Soybean Board. We thank Brian Diers of University of Illinois at Urbana-Champaign for providing characterized soybean accessions, undergraduate John Stuntebeck for providing experiment support, and Nicholas Keuler of the UW-Madison Statistical Consulting Group for advice on statistical analyses.

---

**Supplemental Figure 1.**
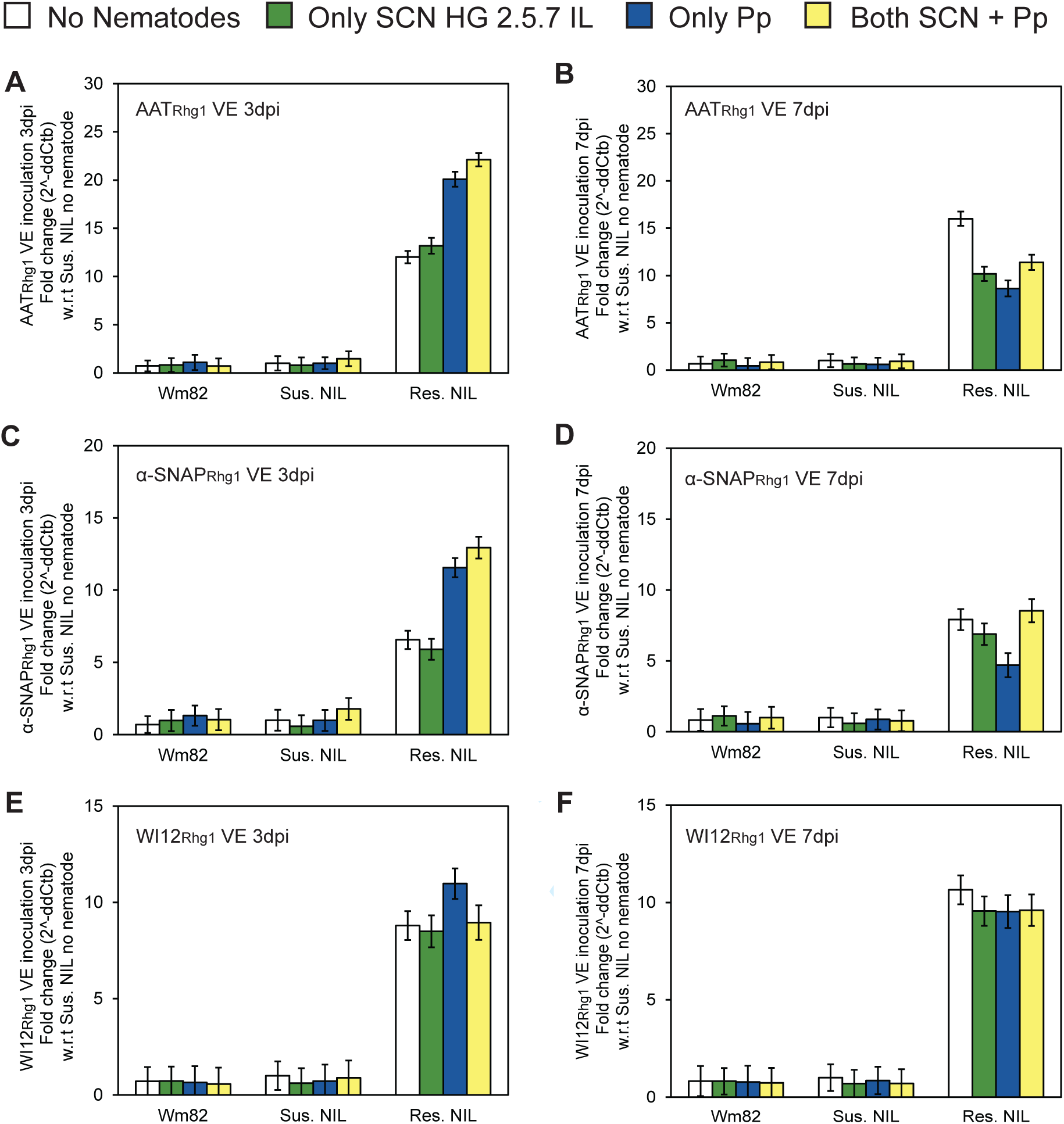
Same data as in Figure 4 but with data for all genotypes normalized to mean Sus. NIL no nematode control within same experiment, to show relative abundances between genotypes. *Rhg1* transcript induction (fold change) compared to no-nematode controls within each plant genotype. The protein encoded by the analyzed transcript is noted in the upper-left corner of each graph. Wm82, Sus. NIL and Res. NIL plants were inoculated at VE stage and root samples from infection areas taken at: (A, C, E and G) 3 dpi, or (B, D, F and H) 7 dpi. No statistically significant differences between samples were observed using ANOVA and Tukey HSD test. The SCN HG 2.5.7 IL used partially overcomes *rhg1-b* resistance and represents the most prevalent HG type in the USA. Data are for two independent experiments with three replicate roots assessed within each experiment (n=6 for each bar).

**Supplemental Figure 2.**
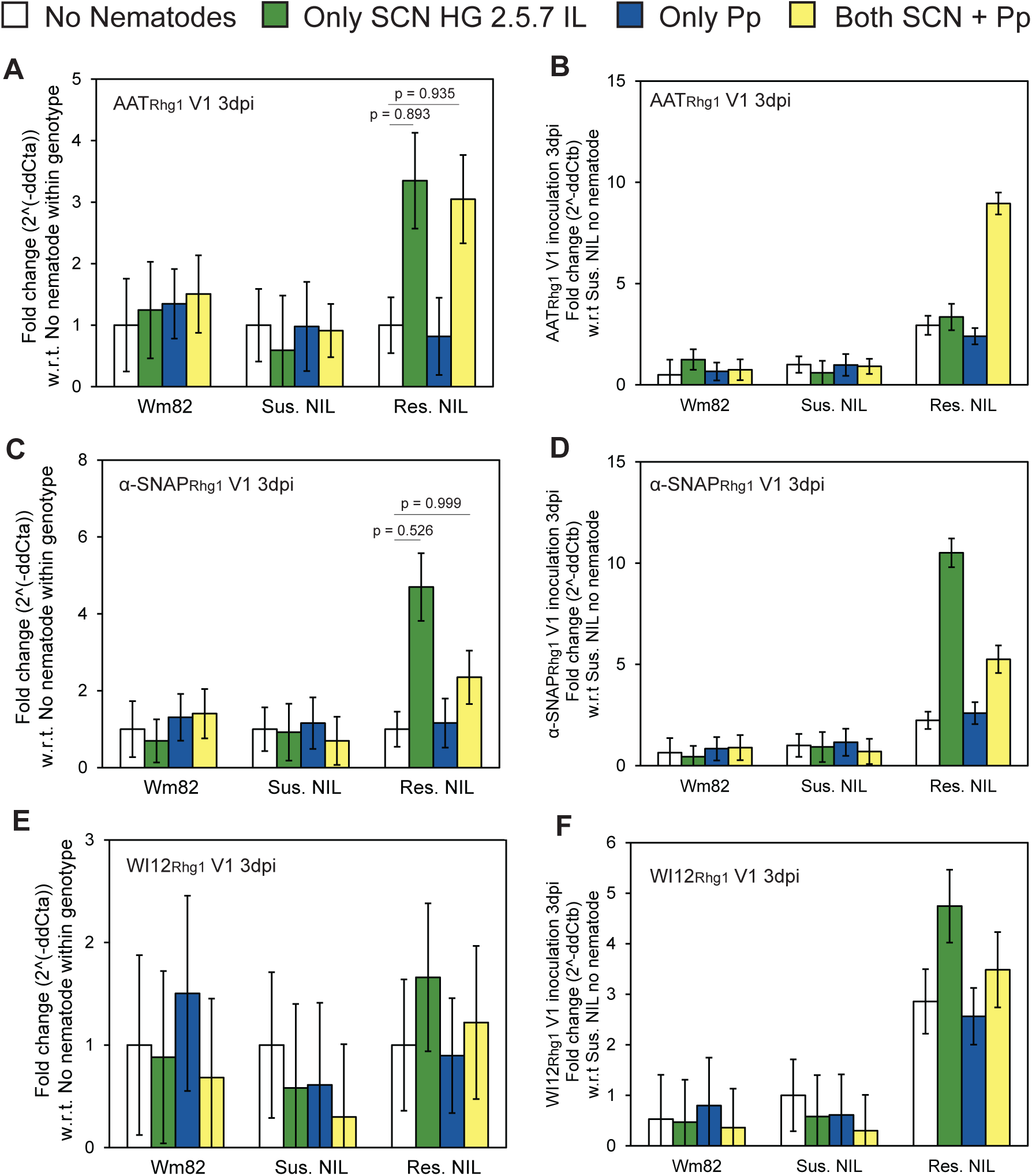
*Rhg1* transcript induction (fold change) for V1 plants at 3 dpi compared to no-nematode controls within each plant genotype. The protein encoded by the analyzed transcript is noted in the upper-left corner of each graph. Wm82, Sus. NIL and Res. NIL plants were inoculated at V1 stage. (A, C, E) Data normalized to mean for no-nematode control of same genotype in same experiment. (B, D, F) Same data as A, C, E with data for all genotypes normalized to mean Sus. NIL no nematode control within same experiment, to show relative abundances between genotypes. Instances of possible significant differences of means are marked with p-values from ANOVA Tukey HSD tests (p > 0.2 if no p value is shown). The SCN HG 2.5.7 IL used partially overcomes *rhg1-b* resistance and represents the most prevalent HG type in the USA. All experiments were repeated twice with two replicate roots assessed within each experiment (n=4 for each bar).

**Supplementary Table 1.**
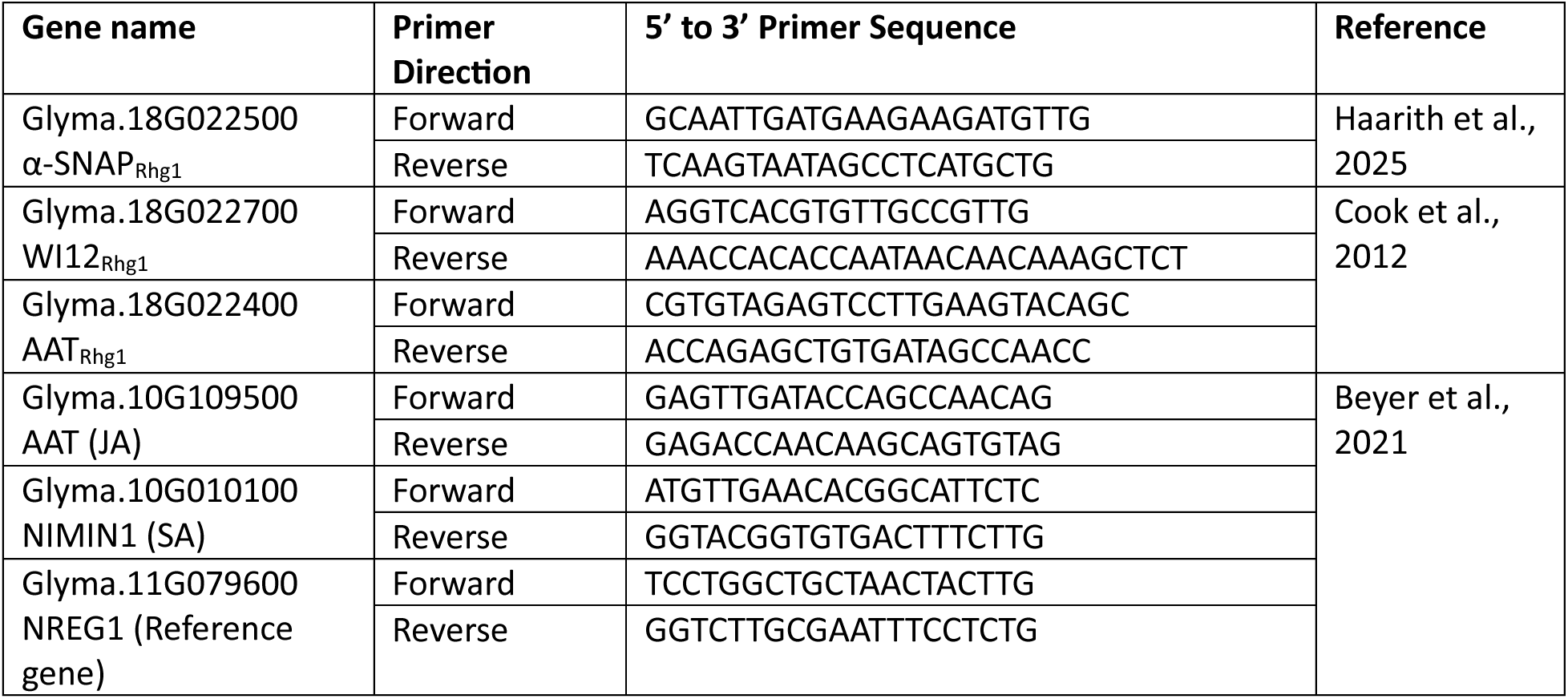
List of all qPCR primers used in this study to quan7fy mRNA/transcript abundance.

**Supplementary Table 2.**
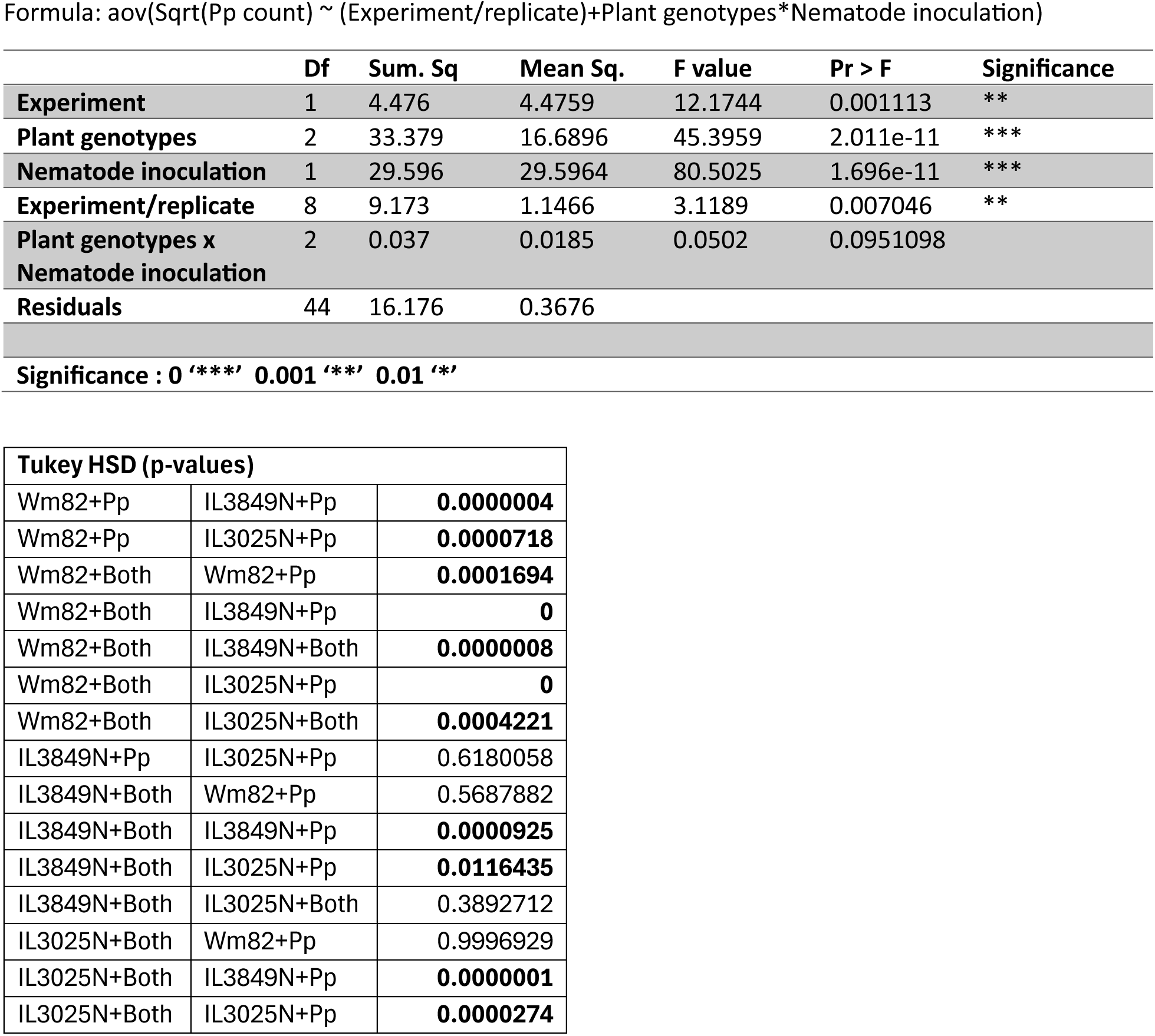
ANOVA for Pp numbers in 1:1 SCN:Pp inocula7on (Wm82, IL3045N and IL3849N) Formula: aov(Sqrt(Pp count) ∼ (Experiment/replicate)+Plant genotypes*Nematode inocula7on)

**Supplementary Table 3.**
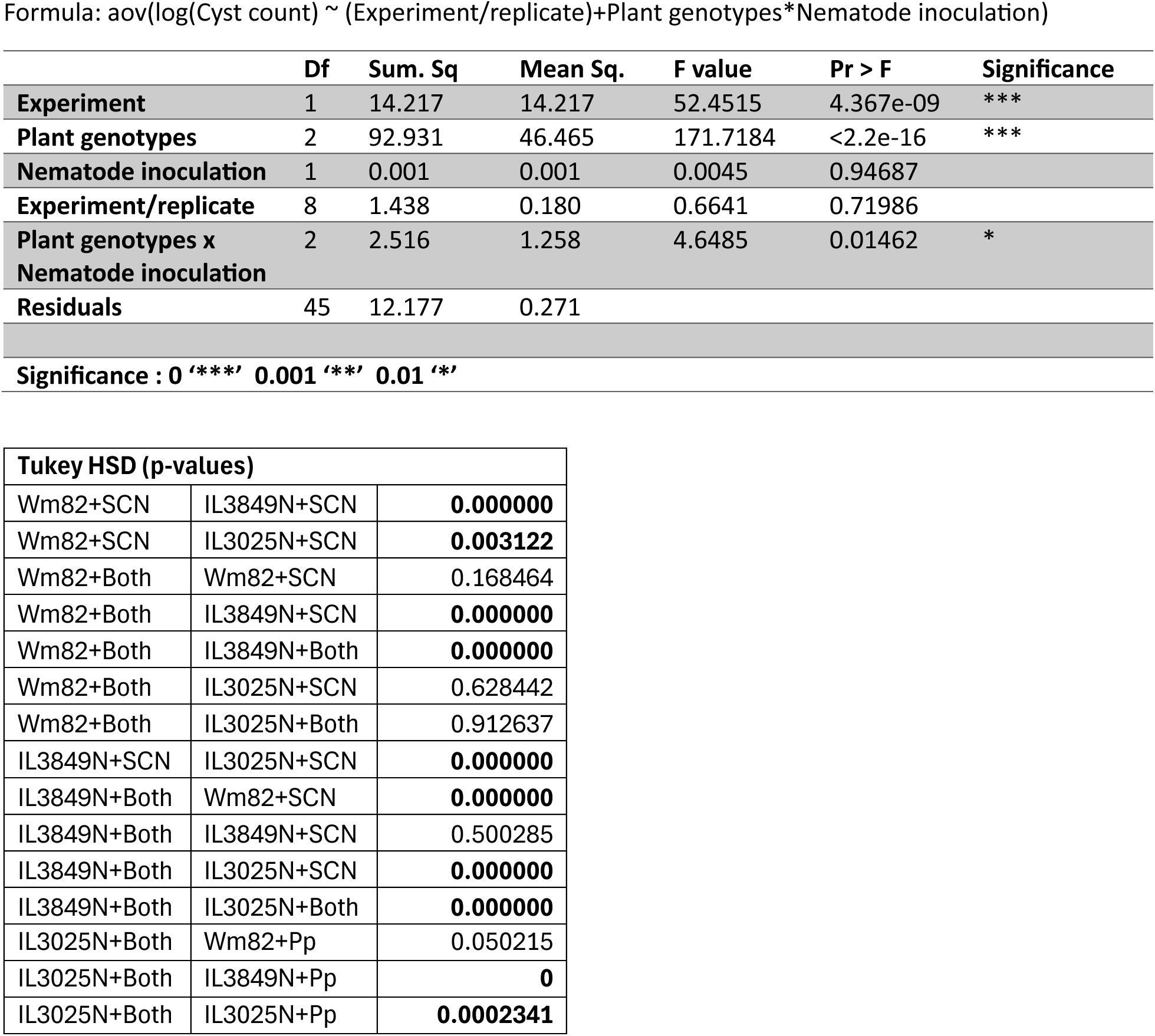
ANOVA for SCN cyst numbers in 1:1 SCN:Pp inocula7on (Wm82, IL3045N and IL3849N) Formula: aov(log(Cyst count) ∼ (Experiment/replicate)+Plant genotypes*Nematode inocula7on)

**Supplementary Table 4.**
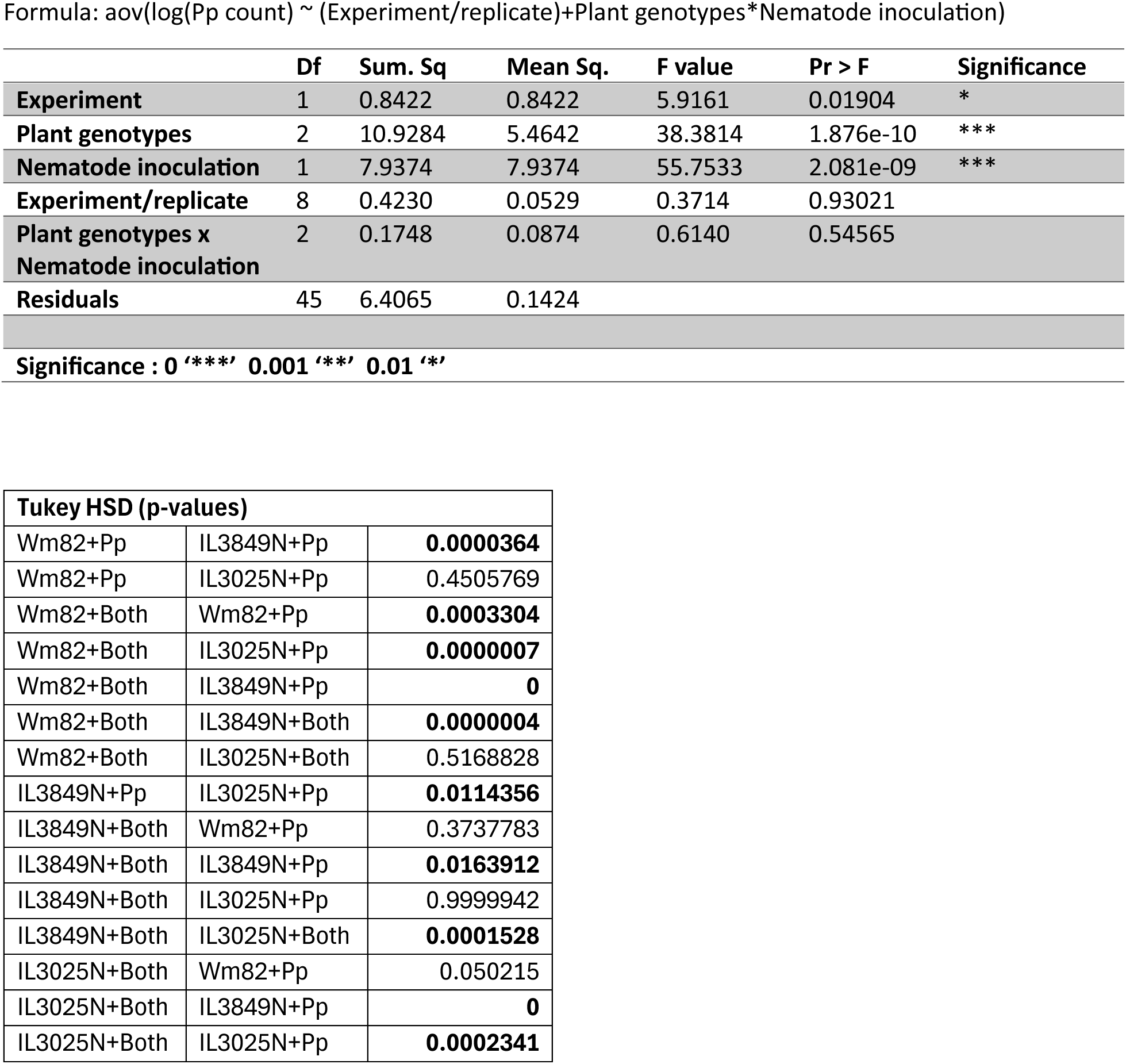
ANOVA for Pp numbers in 2:1 SCN:Pp inocula7on(Wm82, IL3045N and IL3849N) Formula: aov(log(Pp count) ∼ (Experiment/replicate)+Plant genotypes*Nematode inocula7on)

**Supplementary Table 5.**
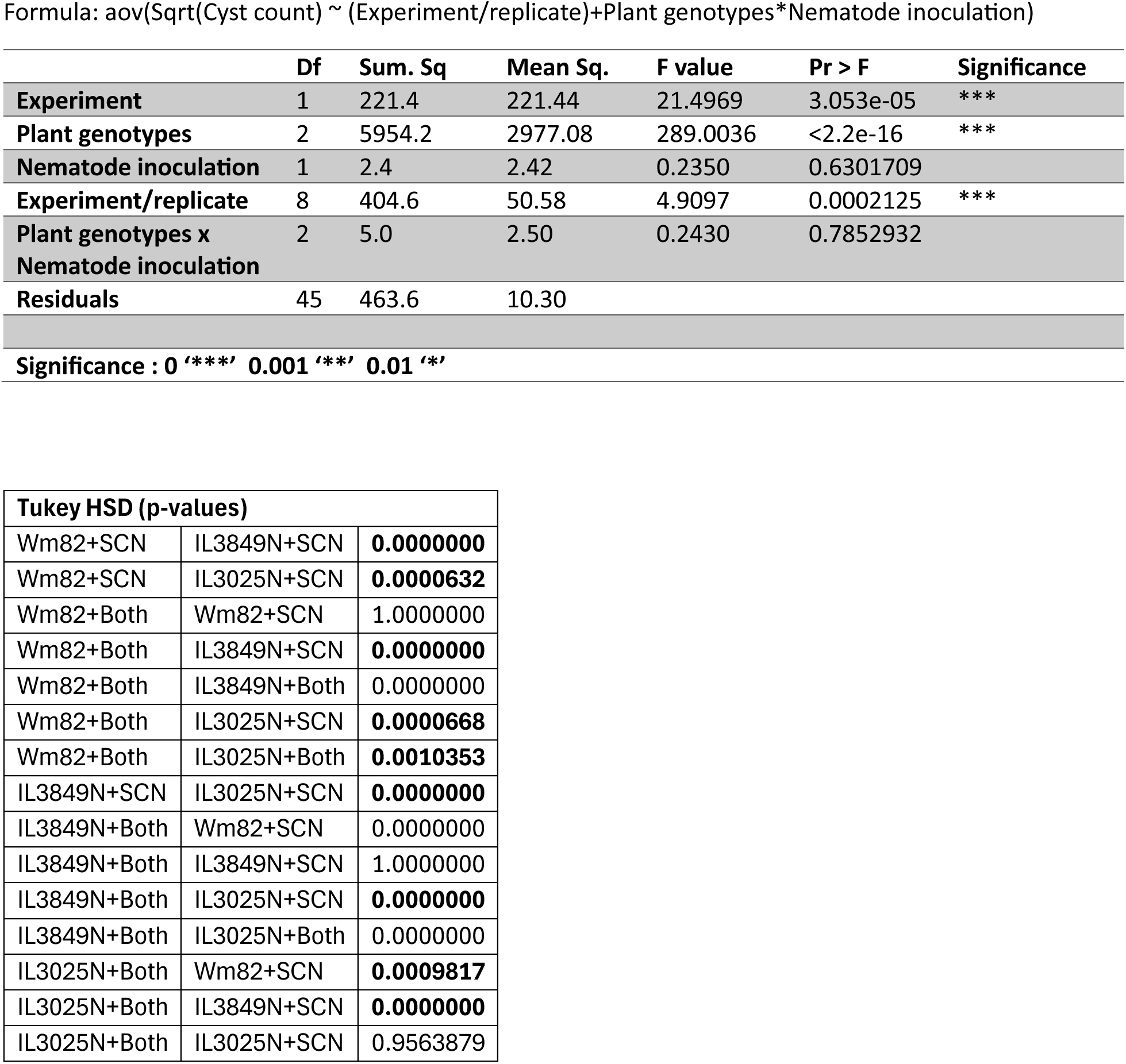
ANOVA for SCN cyst numbers in 2:1 SCN:Pp inocula7on (Wm82, IL3045N and IL3849N) Formula: aov(Sqrt(Cyst count) ∼ (Experiment/replicate)+Plant genotypes*Nematode inocula7on)

**Supplementary Table 6.**
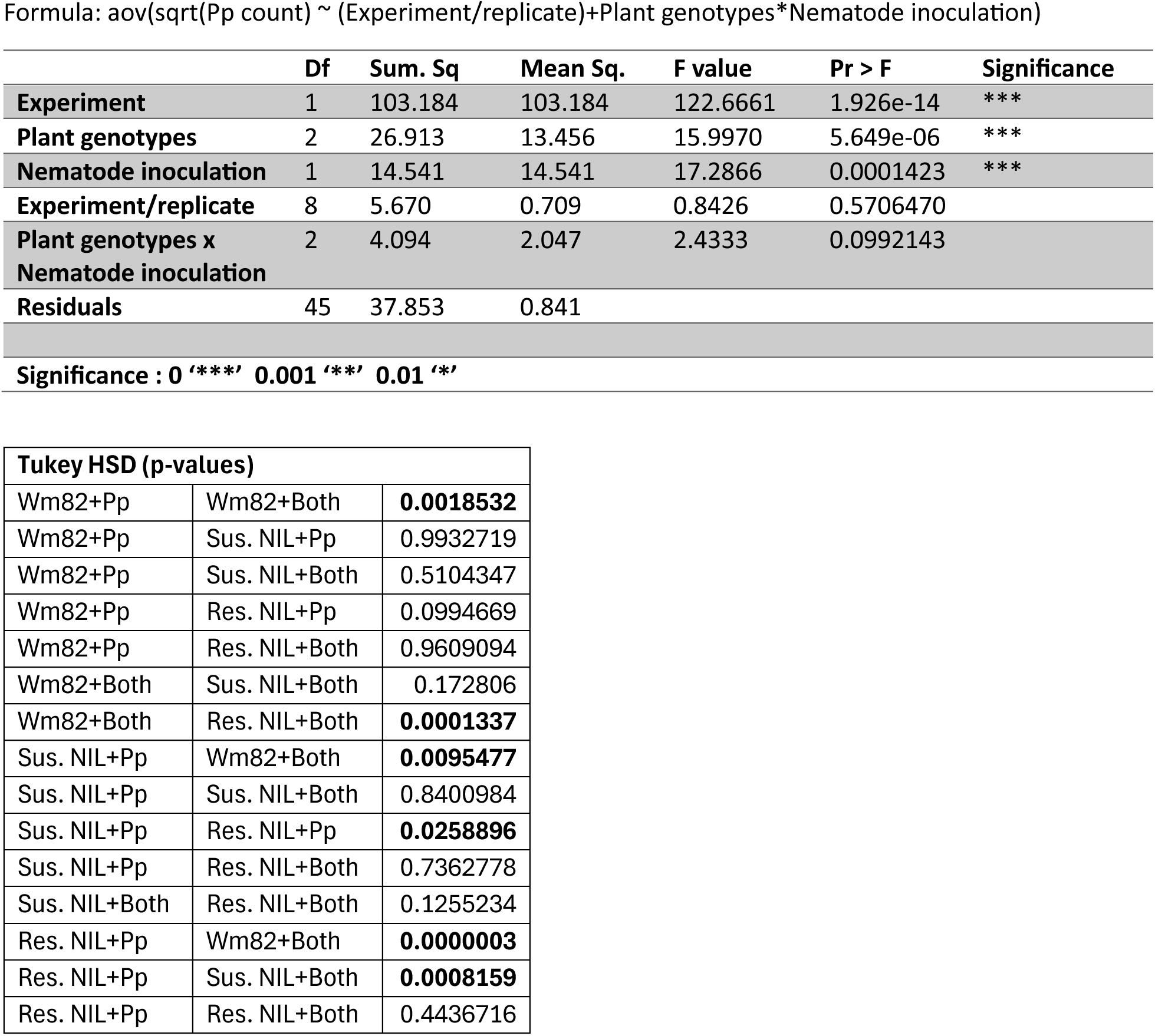
ANOVA for Total Pp numbers on NILs. Formula: aov(sqrt(Pp count) ∼ (Experiment/replicate)+Plant genotypes*Nematode inocula7on)

**Supplementary Table 7.**
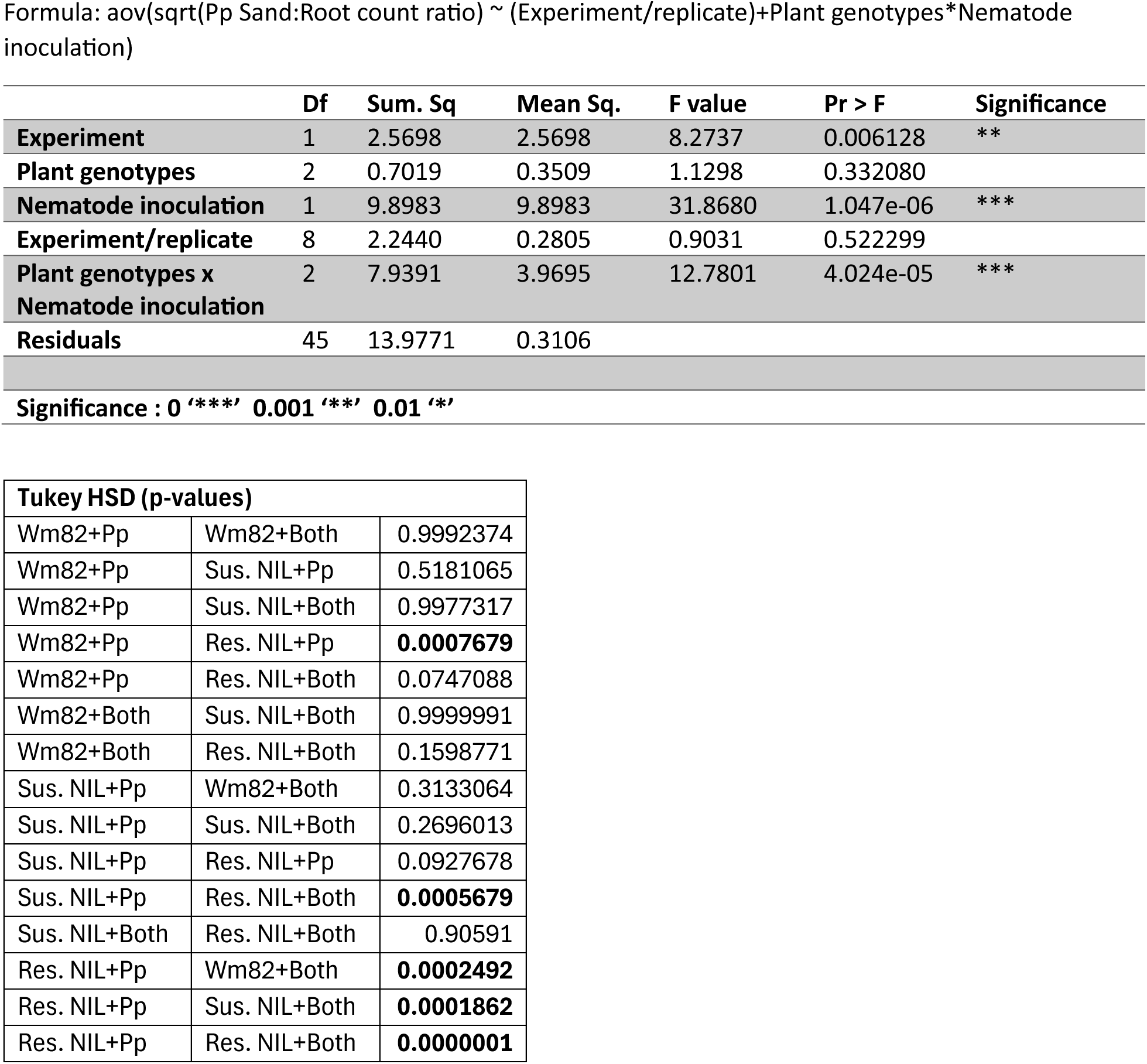
ANOVA for Total Pp Sand:Root ra7o on NILs. Formula: aov(sqrt(Pp Sand:Root count ra7o) ∼ (Experiment/replicate)+Plant genotypes*Nematode inocula7on)

**Supplementary Table 8.**
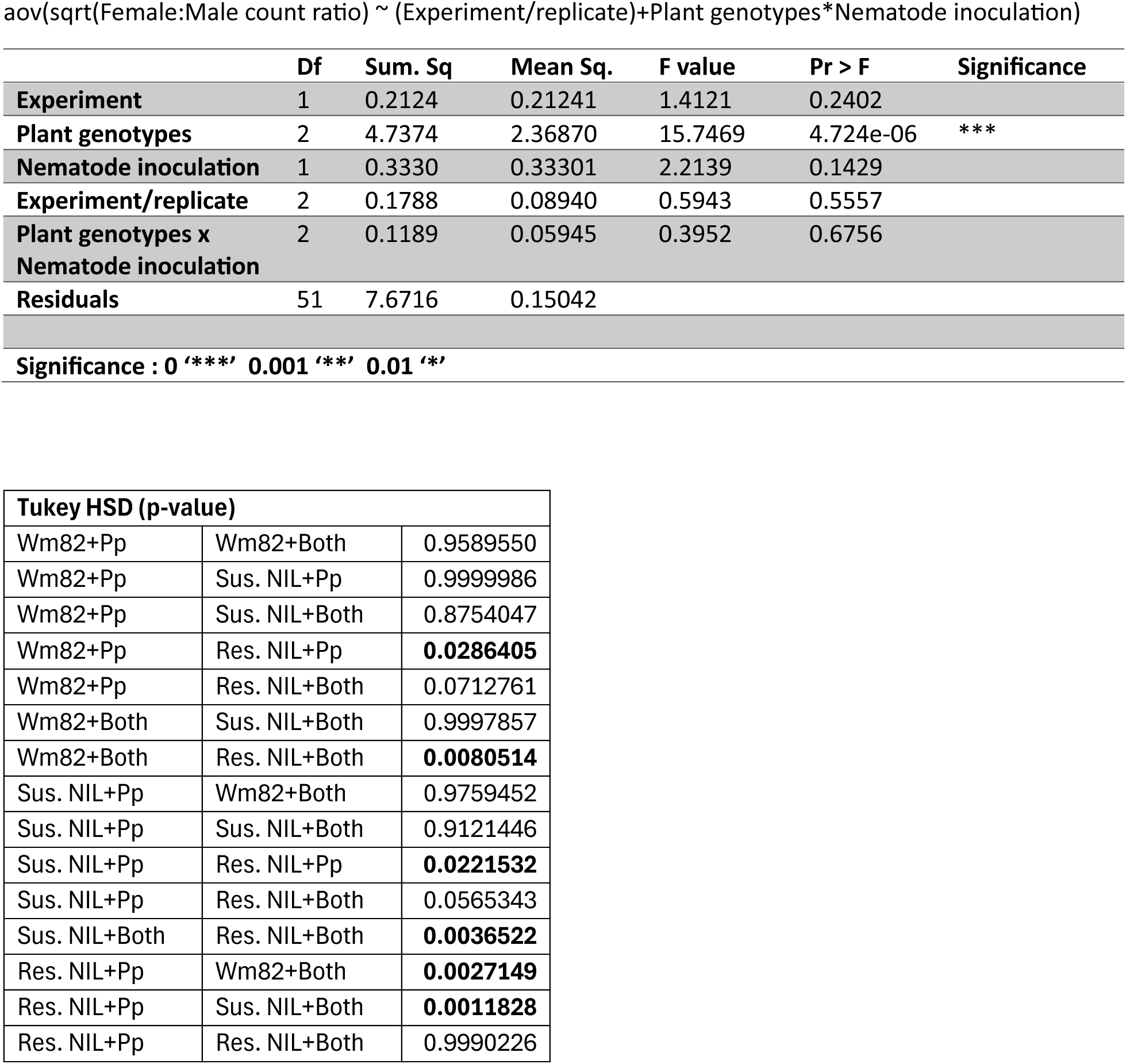
ANOVA for Total Pp Female:Male ra7o on NILs. aov(sqrt(Female:Male count ra7o) ∼ (Experiment/replicate)+Plant genotypes*Nematode inocula7on)

**Supplementary Table 9.**
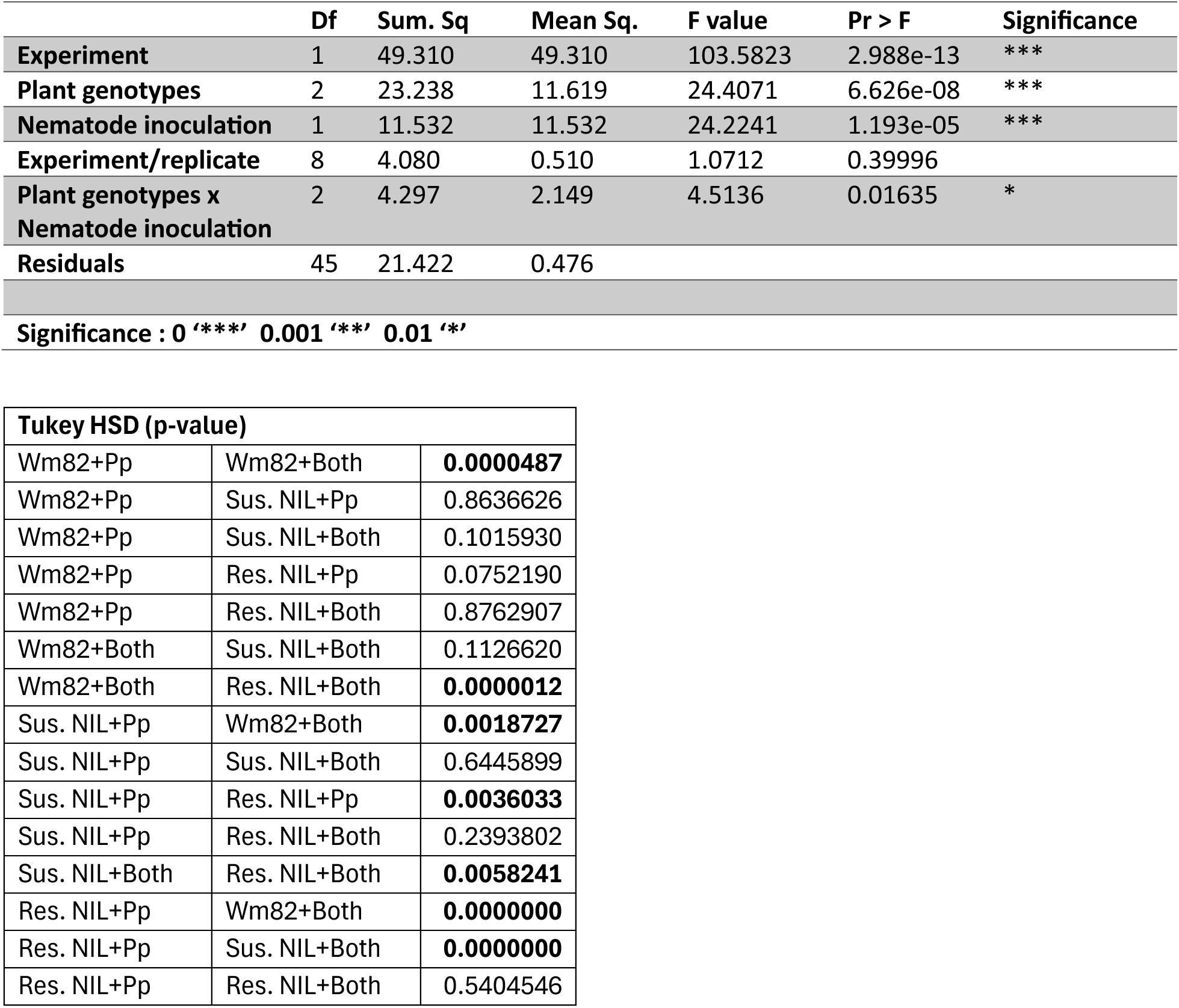
ANOVA for Total Pp Juveniles (J2+J3+J4) on NILs.

**Supplementary Table 10.**
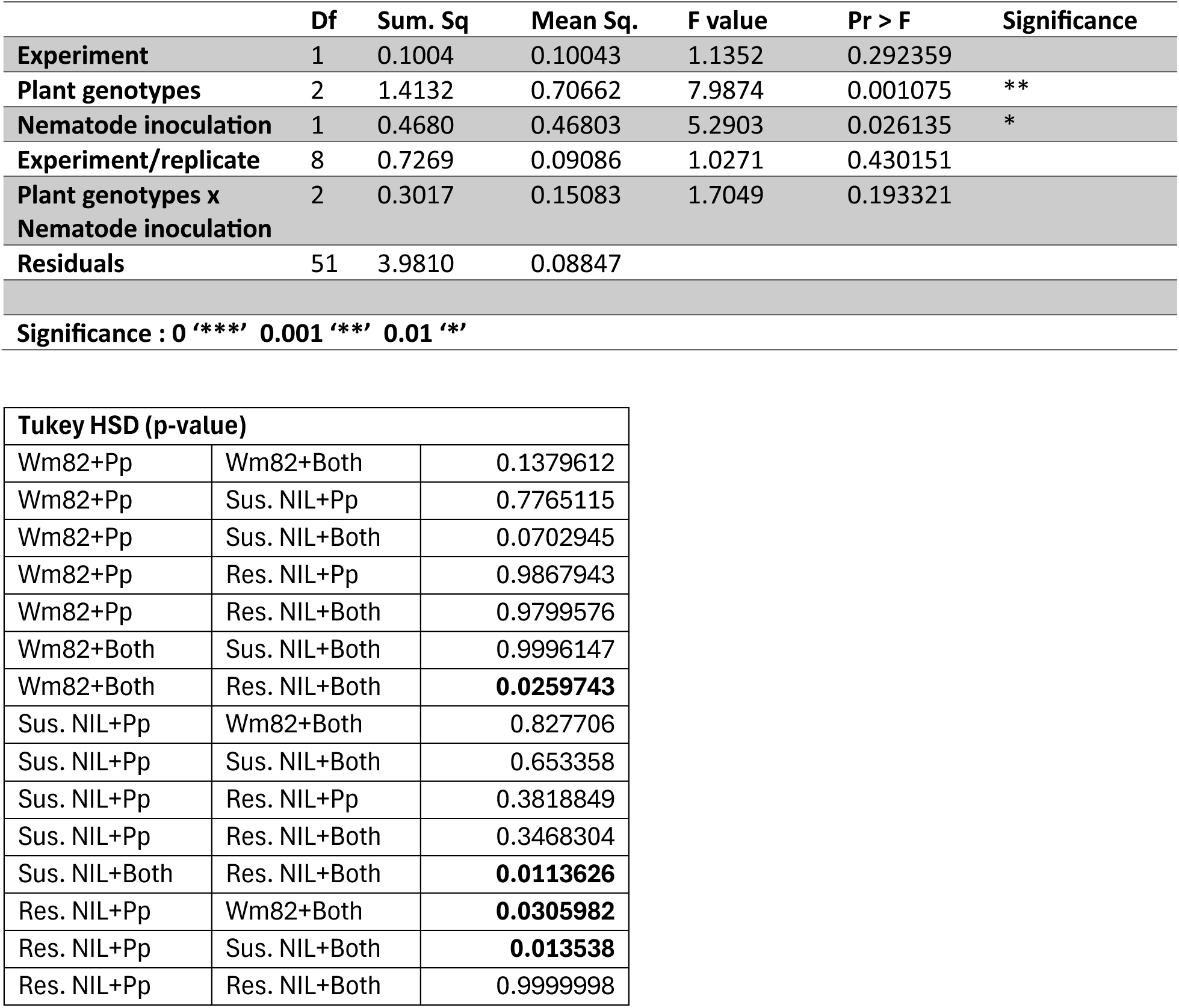
ANOVA for Total Pp Juveniles (J2+J3+J4):Adults (Male+Female) on NILs.

**Supplementary Table 11.**
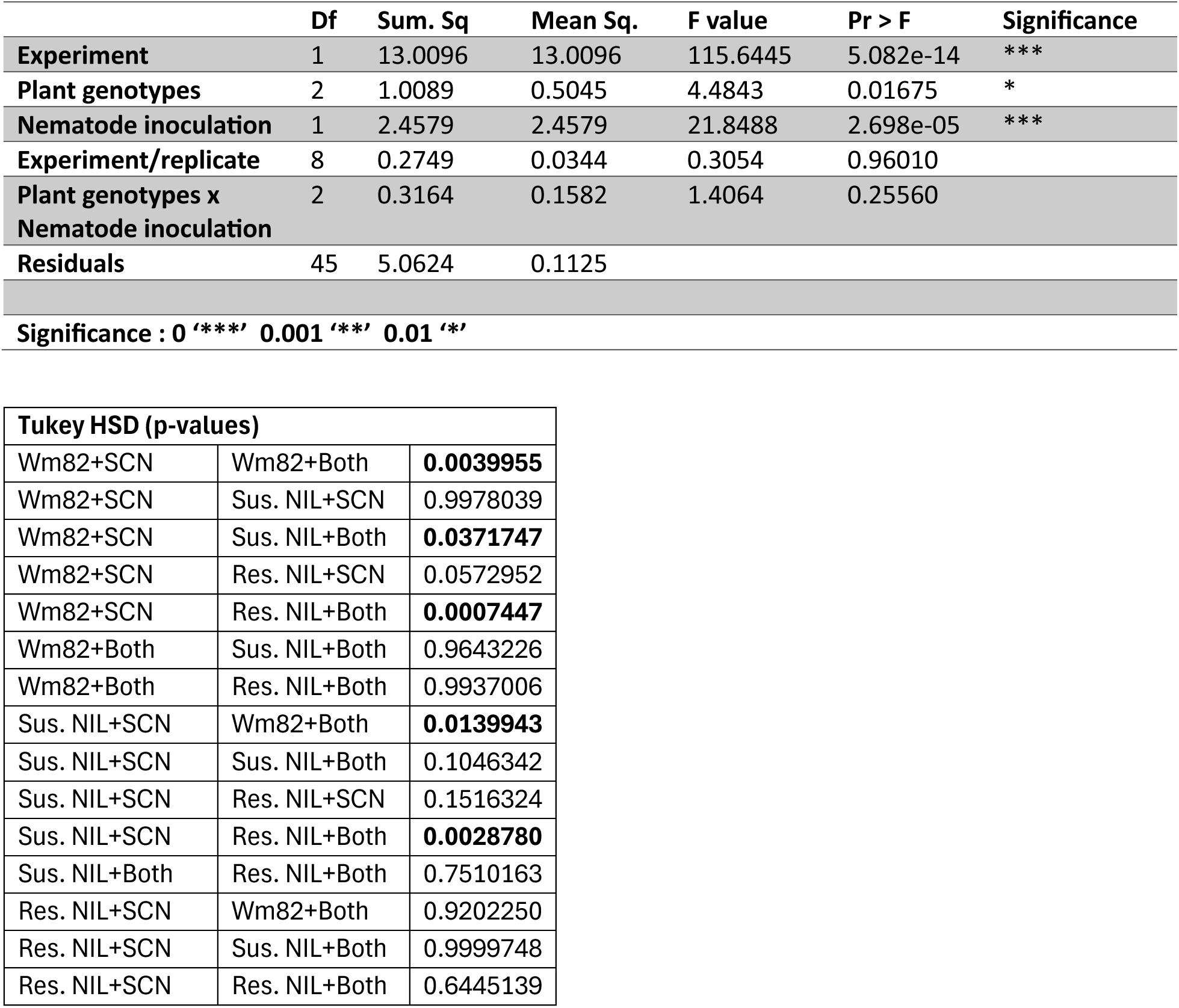
ANOVA for Total SCN cysts on NILs.

**Supplementary Table 12.**
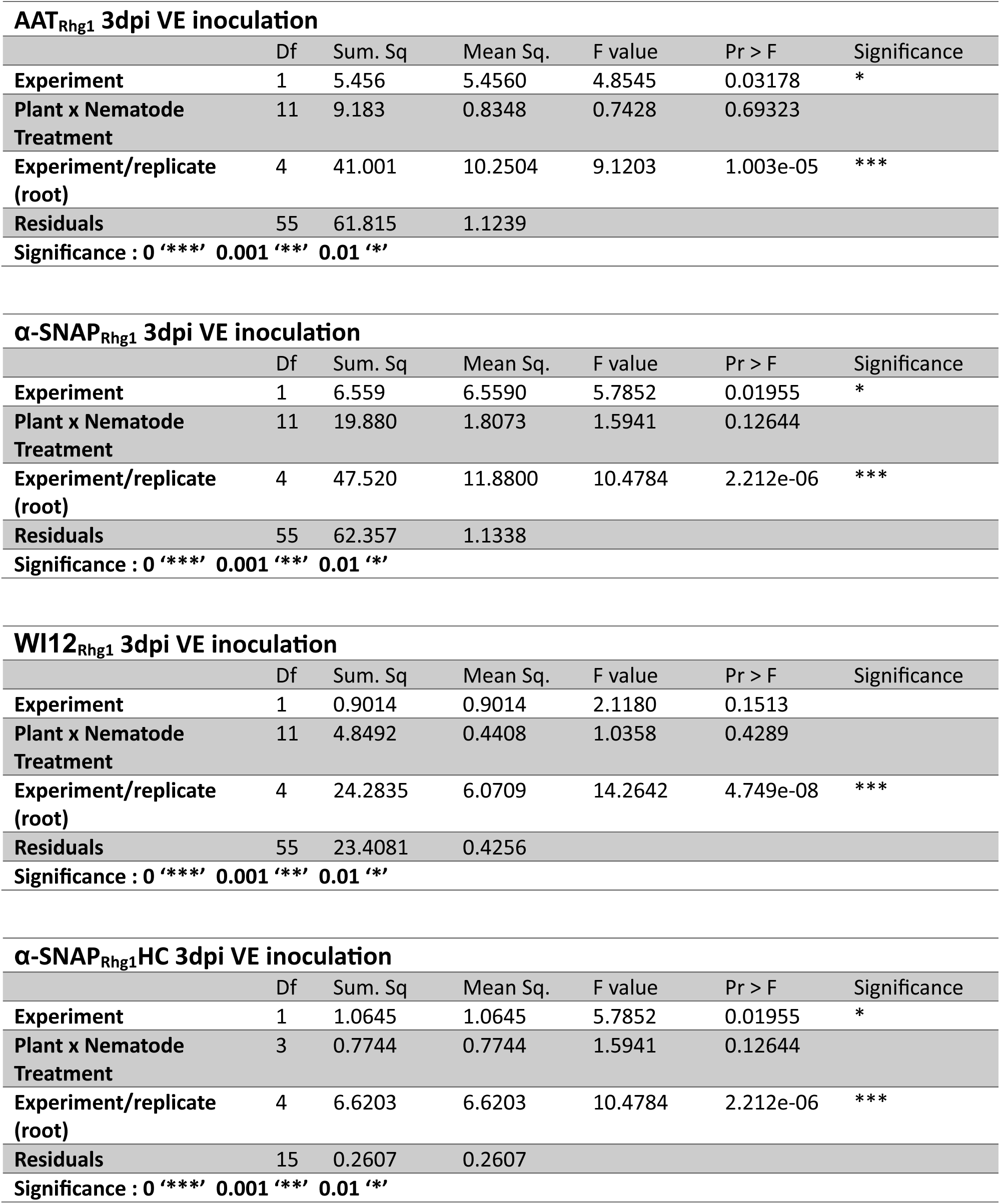
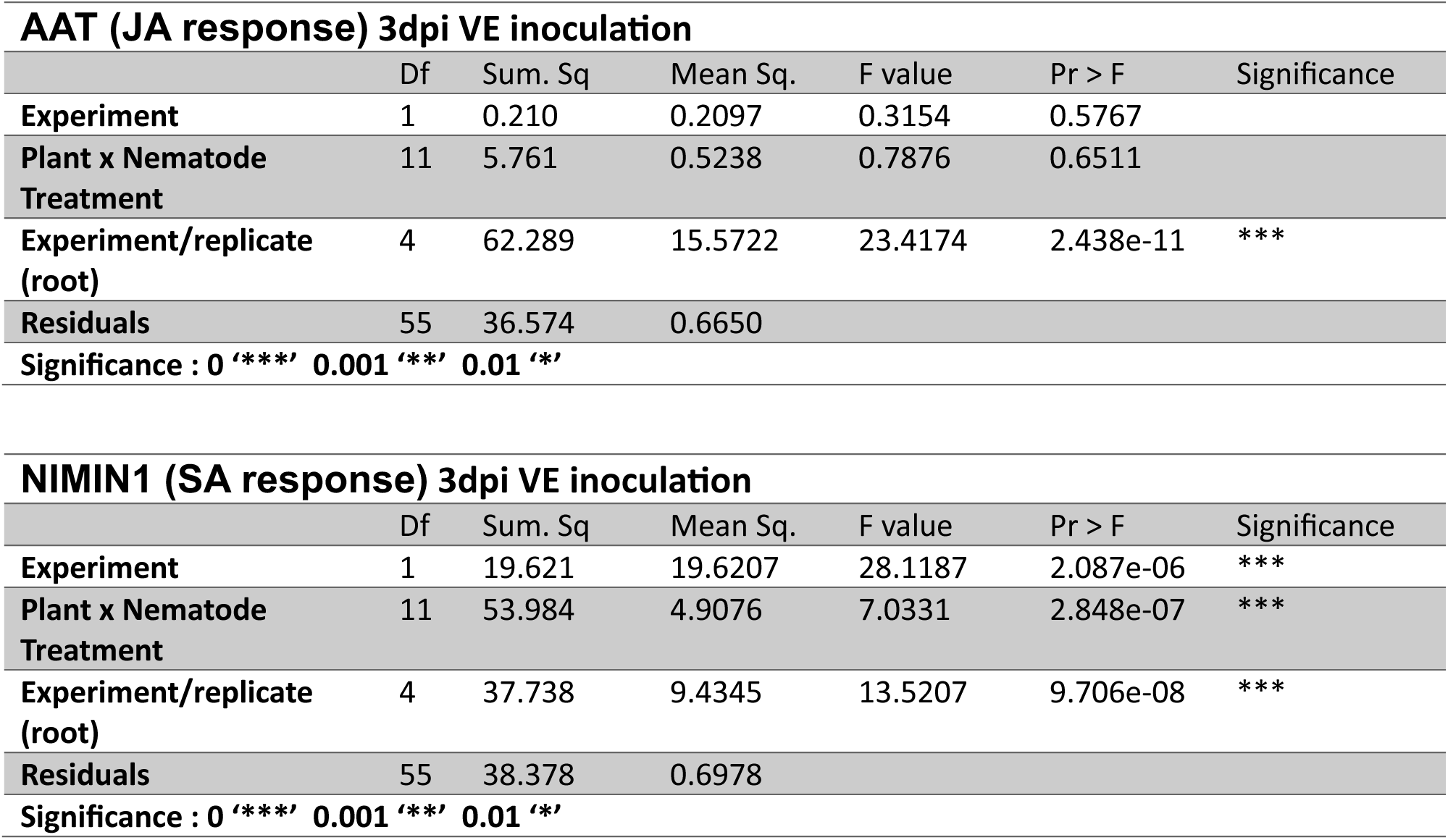
ANOVA for RT q-PCR transcript abundance for 3dpi VE inocula7on 1:1 SCN:Pp. Analyses were done on ddCt values = (Ct sample – CtNREG1) - avg(Ct No nematode sample of the same plant genotype – CtNREG1).

**Supplementary Table 13.**
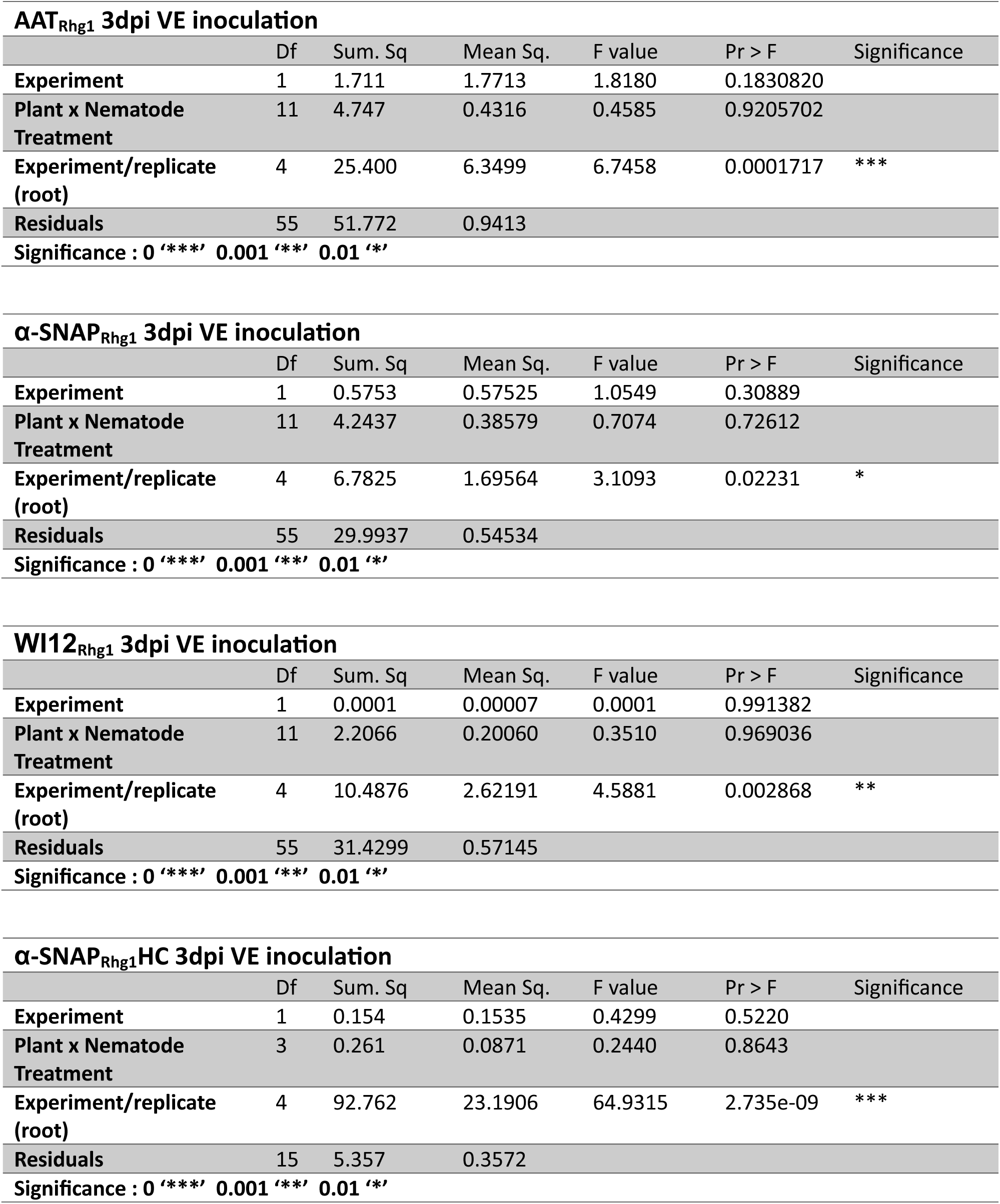
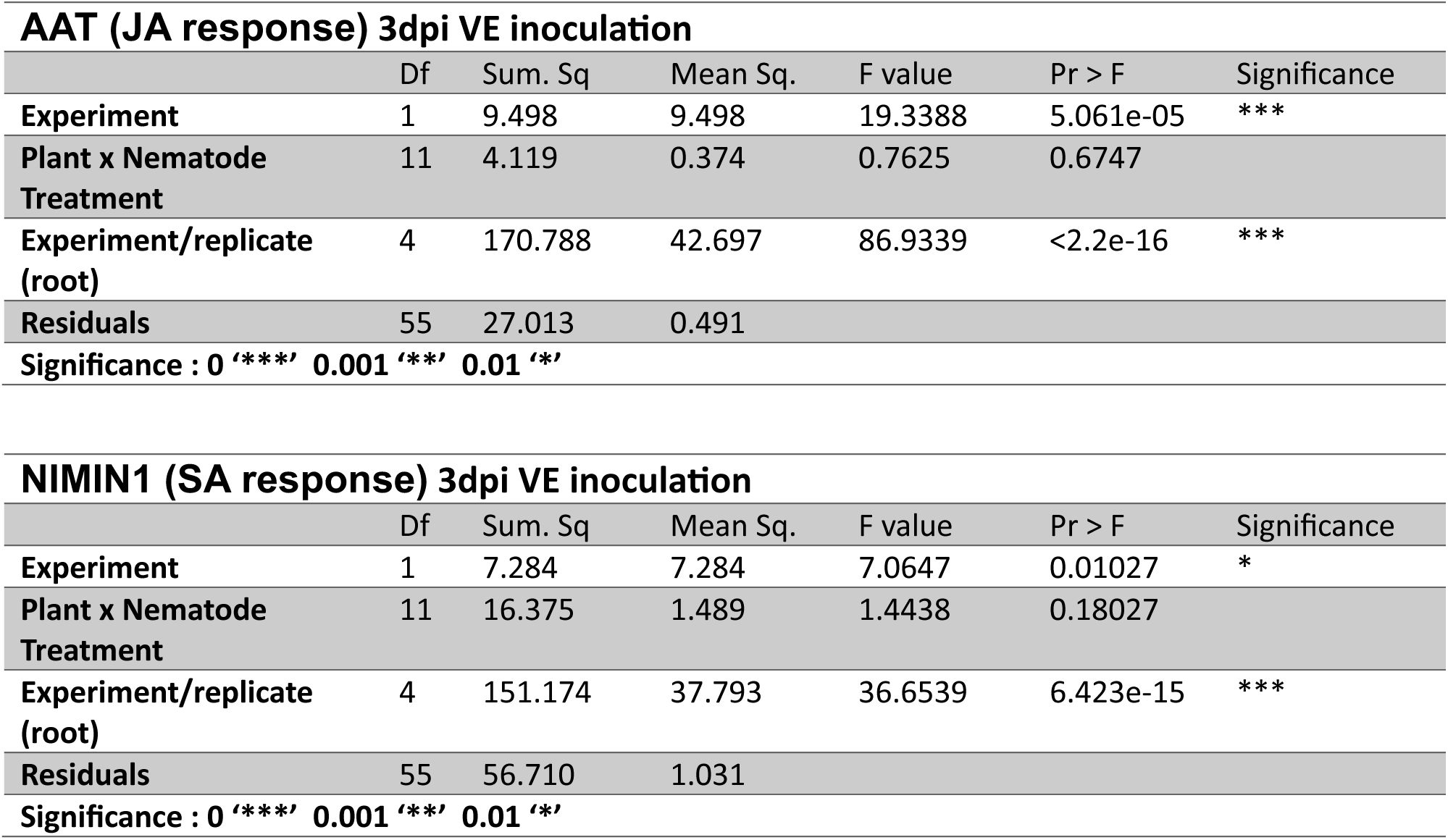
ANOVA for RT q-PCR transcript abundance for 7dpi VE inocula7on 1:1 SCN:Pp. Analyses were done on ddCt values = (Ct sample – CtNREG1) - avg(Ct No nematode sample of the same plant genotype – CtNREG1).

**Supplementary Table 14.**
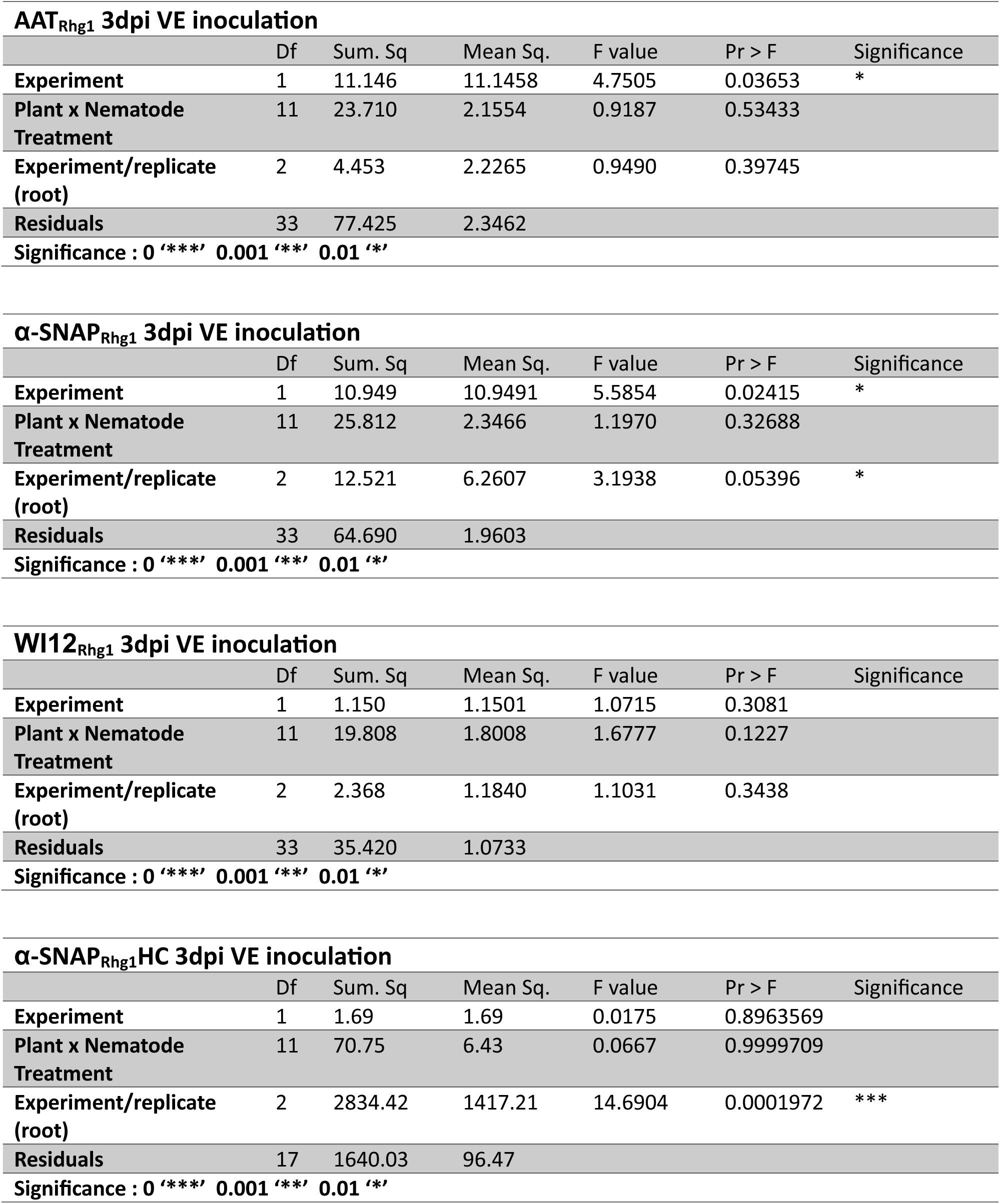
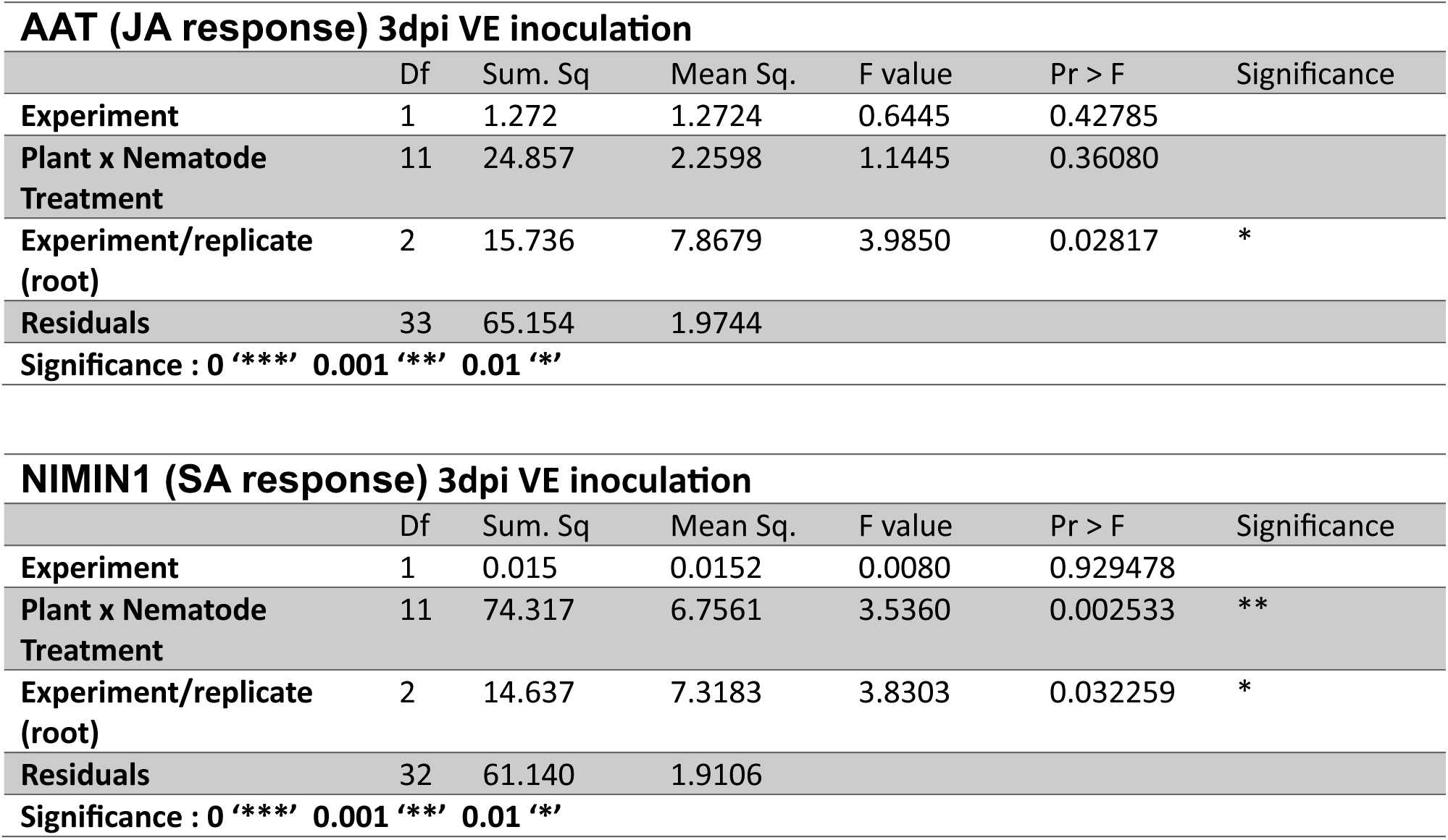
ANOVA for RT q-PCR transcript abundance for 3dpi V1 inocula7on 1:1 SCN:Pp. Analyses were done on ddCt values = (Ct sample – CtNREG1) - avg(Ct No nematode sample of the same plant genotype – CtNREG1).

## Notes

Funding: Wisconsin Soybean Marketing Board award #MSN274578 2023-2024 and United Soybean Board award #25-209-S-C-3-A

### Competing Interest Statement

The authors have declared no competing interest.

### Summary of Updates

The manuscript was revised based on reviewers' comments from Plant Disease journal and was resubmitted on the 2nd of Feb 2026.

## Literature Cited

Allen, T. W., Bradley, C. A., Sisson, A. J., Byamukama, E., Chilvers, M. I., Coker, C. M., Collins, A. A., Damicone, J. P., Dorrance, A. E., Dufault, N. S., Esker, P. D., Faske, T. R., Giesler, L. J., Grybauskas, A. P., Hershman, D. E., Hollier, C. A., Isakeit, T., Jardine, D. J., Kelly, H. M., Kemerait, R. C., Kleczewski, N. M., Koenning, S. R., Kurle, J. E., Malvick, D. K., Markell, S. G., Mehl, H. L., Mueller, D. S., Mueller, J. D., Mulrooney, R. P., Nelson, B. D., Newman, M. A., Osborne, L., Overstreet, C., Padgett, G. B., Phipps, P. M., Price, P. P., Sikora, E. J., Smith, D. L., Spurlock, T. N., Tande, C. A., Tenuta, A. U., Wise, K. A., and Wrather, J. A. 2017. Soybean Yield Loss Estimates Due to Diseases in the United States and Ontario, Canada, from 2010 to 2014. Plant Health Prog. 18:19–27.

Baermann, G. 1917. Eine einfache Methode zur Auffindung von Anklostomum (Nematoden) Larven in Erdproben. Geneeskundig tijdschrift voor Nederlandsch-Indië 57:131–137.

Basnet, P., Meinhardt, C. G., Usovsky, M., Gillman, J. D., Joshi, T., Song, Q., Diers, B., Mitchum, M. G., and Scaboo, A. M. 2022. Epistatic interaction between *Rhg1-a* and *Rhg2* in PI 90763 confers resistance to virulent soybean cyst nematode populations. Theor. Appl. Genet. 135:2025–2039.

Bayless, A. M., Zapotocny, R. W., Grunwald, D. J., Amundson, K. K., Diers, B. W., and Bent, A. F. 2018. An atypical N-ethylmaleimide sensitive factor enables the viability of nematode-resistant *Rhg1* soybeans. Proc. Natl. Acad. Sci. U S A 115:E4512–E4521.

Bayless, A. M., Zapotocny, R. W., Han, S., Grunwald, D. J., Amundson, K. K., and Bent, A. F. 2019. The *rhg1-a* (*Rhg1* low-copy) nematode resistance source harbors a copia-family retrotransposon within the *Rhg1*-encoded alpha-SNAP gene. Plant Direct 3:e00164.

Bayless, A. M., Smith, J. M., Song, J., McMinn, P. H., Teillet, A., August, B. K., and Bent, A. F. 2016. Disease resistance through impairment of alpha-SNAP-NSF interaction and vesicular trafficking by soybean *Rhg1*. Proc. Natl. Acad. Sci. U S A 113:E7375–E7382.

Bent, A. 2022. Exploring Soybean Resistance to Soybean Cyst Nematode. Annu. Rev. Phytopathol. 60:379–409.

Beyer, S. F., Bel, P. S., Flors, V., Schultheiss, H., Conrath, U., and Langenbach, C. J. G. 2021. Disclosure of salicylic acid and jasmonic acid-responsive genes provides a molecular tool for deciphering stress responses in soybean. Sci. Rep. 11:20600.

Bradley, C. A., Allen, T. W., Sisson, A. J., Bergstrom, G. C., Bissonnette, K. M., Bond, J., Byamukama, E., Chilvers, M. I., Collins, A. A., Damicone, J. P., Dorrance, A. E., Dufault, N. S., Esker, P. D., Faske, T. R., Fiorellino, N. M., Giesler, L. J., Hartman, G. L., Hollier, C. A., Isakeit, T., Jackson-Ziems, T. A., Jardine, D. J., Kelly, H. M., Kemerait, R. C., Kleczewski, N. M., Koehler, A. M., Kratochvil, R. J., Kurle, J. E., Malvick, D. K., Markell, S. G., Mathew, F. M., Mehl, H. L., Mehl, K. M., Mueller, D. S., Mueller, J. D., Nelson, B. D., Overstreet, C., Padgett, G. B., Price, P. P., Sikora, E. J., Small, I., Smith, D. L., Spurlock, T. N., Tande, C. A., Telenko, D. E. P., Tenuta, A. U., Thiessen, L. D., Warner, F., Wiebold, W. J., and Wise, K. A. 2021. Soybean Yield Loss Estimates Due to Diseases in the United States and Ontario, Canada, from 2015 to 2019. Plant Health Prog. 22:483–495.

Brucker, E., Carlson, S., Wright, E., Niblack, T., and Diers, B. 2005. *Rhg1* alleles from soybean PI 437654 and PI 88788 respond differentially to isolates of Heterodera glycines in the greenhouse. Theor. Appl. Genet. 111:44–49.

Brzostowski, L. F., and Diers, B. W. 2017. Pyramiding of Alleles from Multiple Sources Increases the Resistance of Soybean to Highly Virulent Soybean Cyst Nematode Isolates. Crop. Sci. 57:2932–2941.

Butler, K. J., Chen, S., Smith, J. M., Wang, X., and Bent, A. F. 2019. Soybean Resistance Locus *Rhg1* Confers Resistance to Multiple Cyst Nematodes in Diverse Plant Species. Phytopathol. 109:2107–2115.

Butler, K. J., Fliege, C., Zapotocny, R., Diers, B., Hudson, M., and Bent, A. F. 2021. Soybean Cyst Nematode Resistance Quantitative Trait Locus *cqSCN-006* Alters the Expression of a gamma-SNAP Protein. Mol. Plant Microbe Interact. 34:1433–1445.

Castanheira, C. M., Falcão, H. G., Ida, E. I., Dias-Arieira, C. R., and Bolanho Barros, B. C. 2020. *Pratylenchus brachyurus* parasitism on soybean: effects on productivity, vegetative and nematological parameters and chemical properties. Eur. J. Plant Pathol. 157:651–661.

Chen, S., Shaeffer, C. C., Wyse, D. L., Nickel, P., and Kandel, H. 2012. Plant-parasitic Nematode Communities and Their Associations with Soil Factors in Organically Farmed Fields in Minnesota. J. Nematol. 44:361–369.

Chowdhury, I. A., Yan, G. P., Kandel, H., and Plaisance, A. 2022. Population Development of the Root-Lesion Nematode on Soybean Cultivars. Plant Dis. 106:2117–2126.

Colgrove, A. L., and Niblack, T. L. 2005. The effect of resistant soybean on male and female development and adult sex ratios of. J. Nematol. 37:161–167.

Cook, D. E., Bayless, A. M., Wang, K., Guo, X., Song, Q., Jiang, J., and Bent, A. F. 2014. Distinct Copy Number, Coding Sequence, and Locus Methylation Patterns Underlie *Rhg1*-Mediated Soybean Resistance to Soybean Cyst Nematode. Plant Physiol. 165:630–647.

Cook, D. E., Lee, T. G., Guo, X., Melito, S., Wang, K., Bayless, A. M., Wang, J., Hughes, T. J., Willis, D. K., Clemente, T. E., Diers, B., Jiang, J., Hudson, M., and Bent, A. F. 2012. Copy Number Variation of Multiple Genes at *Rhg1* Mediates Nematode Resistance in Soybean. Science 338:1206–1209.

Dong, J., and Hudson, M. E. 2022. WI12(Rhg1) interacts with DELLAs and mediates soybean cyst nematode resistance through hormone pathways. Plant Biotechnol. J. 20:283–296.

Endo, B. Y. 1967. Comparative Population Increase of *Pratylenchus brachyurus* and *P. zeae* in Corn and in Soybean Varieties Lee and Peking. Phytopathol. 57:118–120.

Favoreto, L., Meyer, M. C., Dias-Arieira, C. R., Machado, A. C. Z., Santiago, D. C., and Riberiro, N. R. 2019. Diagnosis and Management of Nematodes In Soybean Crop. Informe Agropecuario 40:18–29.

Glazebrook, J., Chen, W. J., Estes, B., Chang, H. S., Nawrath, C., Metraux, J. P., Zhu, T., and Katagiri, F. 2003. Topology of the network integrating salicylate and jasmonate signal transduction derived from global expression phenotyping. Plant J. 34:217–228.

Haarith, D., Das, S., Nelson, E., Zapotocny, R., and Bent, A. F. 2025. Overexpression of α-SNAP_Rhg1_ Can Improve *rhg1-a* Mediated Soybean Resistance to Soybean Cyst Nematode. Phytopathol.115(9): 10.1094/PHYTO-02-25-0077-R

Hamawaki, O. T., Hamawaki, R. L., Nogueira, A. P. O., Glasenapp, J. S., Hamawaki, C. D. L., and da Silva, C. O. 2019. Evaluation of soybean breeding lineages to new sources of root-knot nematode resistance. Cienc. Agrotec. 43. 10.1590/1413-7054201943009519

Han, S., Smith, J. M., Du, Y., and Bent, A. F. 2023. Soybean transporter AAT_Rhg1_ abundance increases along the nematode migration path and impacts vesiculation and ROS. Plant. Physiol. 192:133–153.

Jones, J. T., Haegeman, A., Danchin, E. G., Gaur, H. S., Helder, J., Jones, M. G., Kikuchi, T., Manzanilla-Lopez, R., Palomares-Rius, J. E., Wesemael, W. M., and Perry, R. N. 2013. Top 10 plant-parasitic nematodes in molecular plant pathology. Mol. Plant Pathol. 14:946–961.

Kleczewski, N., Colsgrove, A., Bowman, D., Harbach, C., and Plewa, D. 2023. A Survey of Soybean Cyst Nematode Population Densities and Phenotypes in Illinois: 2018 and 2020. Plant Health Prog. 24:221–225.

Lakhssassi, N., Liu, S., Bekal, S., Zhou, Z., Colantonio, V., Lambert, K., Barakat, A., and Meksem, K. 2017. Characterization of the Soluble NSF Attachment Protein gene family identifies two members involved in additive resistance to a plant pathogen. Sci. Rep. 7:45226.

Lasserre, F., Rivoal, R., and Cook, R. 1994. Interactions between Heterodera avenae and Pratylenchus neglectus on Wheat. J. Nematol. 26:336–344.

Liu, S., Kandoth, P. K., Lakhssassi, N., Kang, J., Colantonio, V., Heinz, R., Yeckel, G., Zhou, Z., Bekal, S., Dapprich, J., Rotter, B., Cianzio, S., Mitchum, M. G., and Meksem, K. 2017. The soybean *GmSNAP18* gene underlies two types of resistance to soybean cyst nematode. Nat. Commun. 8:14822.

Livak, K. J. and Schmittgen, T. D. 2001. Analysis of relative gene expression data using real-time quantitative PCR and the 2^-ΔΔCt method. Methods 25:402–8.

Lopez-Nicora, H. D., Ralston, T. I., Diers, B. W., Dorrance, A. E., and Niblack, T. L. 2023. Interactions Among *Heterodera glycines*, *Macrophomina phaseolina*, and Soybean Genotype. Plant Dis. 107:401–412.

MacGuidwin, A. E., and Bender, B. E. 2012. Estimating Population Densities of Root Lesion Nematodes, *Pratylenchus* spp., from Soil Samples Using Dual Active and Passive Assays. Plant Health Prog. 13.E1535–1025.

MacGuidwin, A. E., and Bender, B. E. 2016. Development of a Damage Function Model for *Pratylenchus penetrans* on Corn. Plant Dis. 100:764–769.

MacGuidwin, A. E., Smith, D. L., Conley, S. P., and Saikai, K. A. 2023. Prevalence of Pest Nematodes Associated with Soybean (*Glycine max*) in Wisconsin from 1998 to 2021. J. Nematol. 55:20230053.

Machado, A. C. Z., and Araúujo, J. V. 2016. Broad-sense heritability and variance component estimates for resistance in Brazilian soybean genotypes. Tropical Plant Pathology 41:390–396.

Matsye, P. D., Lawrence, G. W., Youssef, R. M., Kim, K. H., Lawrence, K. S., Matthews, B. F., and Klink, V. P. 2012. The expression of a naturally occurring, truncated allele of an alpha-SNAP gene suppresses plant parasitic nematode infection. Plant Mol Biol 80:131–155.

McCarville, M. T., Marett, C. C., Mullaney, M. P., Gebhart, G. D., and Tylka, G. L. 2017. Increase in Soybean Cyst Nematode Virulence and Reproduction on Resistant Soybean Varieties in Iowa From 2001 to 2015 and the Effects on Soybean Yields. Plant Health Prog. 18:146–155.

Melakeberhan, H., and Dey, J. 2003. Competition between *Heterodera glycines* and *Meloidogyne incognita* or *Pratylenchus penetrans*: Independent Infection Rate Measurements. J. Nematol. 35:1–6.

Mokrini, F., Viaene, N., Waeyenberge, L., Dababat, A. A., and Moens, M. 2018. Investigation of resistance to *Pratylenchus penetrans* and *P. thornei* in international wheat lines and its durability when inoculated together with the cereal cyst nematode *Heterodera avenae*, using qPCR for nematode quantification. Eur. J. Plant Pathol.151:875–889.

Niblack, T. L., Hussey, R. S., and Boerma, H. R. 1986. Effects of Interactions among *Heterodera glycines*, *Meloidogyne incognita*, and Host Genotype on Soybean Yield and Nematode Population-Densities. J. Nematol. 18:436–443.

Niblack, T. L., Lambert, K. N., and Tylka, G. L. 2006. A model plant pathogen from the kingdom animalia: Heterodera glycines, the soybean cyst nematode. Annu. Rev. Phytopathol. 44:283–303.

Nomura, R. B. G., Lopes-Caitar, V. S., Hishinuma-Silva, S. M., Machado, A. C. Z., Meyer, M. C., and Marcelino-Guimarães, F. C. 2024. *Pratylenchus brachyurus*: status and perspectives in Brazilian agriculture. Trop. Plant Pathol. 49:573–589.

Patil, G. B., Lakhssassi, N., Wan, J., Song, L., Zhou, Z., Klepadlo, M., Vuong, T. D., Stec, A. O., Kahil, S. S., Colantonio, V., Valliyodan, B., Rice, J. H., Piya, S., Hewezi, T., Stupar, R. M., Meksem, K., and Nguyen, H. T. 2019. Whole-genome re-sequencing reveals the impact of the interaction of copy number variants of the *rhg1* and *Rhg4* genes on broad-based resistance to soybean cyst nematode. Plant Biotechnol. J. 17:1595–1611.

Pedro, M. S., da Silva, S. A., Picoli, L. H., da Cunha, L. S., and Machado, A. C. Z. 2024. Experimental and statistical approaches to evaluate the reaction of soybean to Pratylenchus brachyurus. Trop. Plant Pathol. 49:612–621.

Pedersen, P. 2009. Soybean Growth and Development. Iowa State University Extension publication PM1945.

Rebois, R.V. and Huettel, R. N. 1986. Population-Dynamics, Root Penetration, and Feeding-Behavior of Pratylenchus-Agilis in Monoxenic Root Cultures of Corn, Tomato, and Soybean. J. Nematology 18:392–397.

Rosa de Araújo, T., de Assis dos Santos Diniz, F., Mendes, B. L., Almeida, L. C., and Santiago, T. R. 2025. *Pratylenchus zeae* is the main root lesion nematode affecting sugarcane in the world’s most important producing regions. Physiol. Mol. Plant Pathol.136: 102576.

Saikai, K., and MacGuidwin, A. E. 2020. Difference in Lesion Formation by Male and Female *Pratylenchus penetrans*. J. Nematol. 52.E2020–90.

Saikai, K., and MacGuidwin, A. E. 2022. Impact of *Pratylenchus penetrans* on Soybean Grown in Wisconsin, U.S.A. Plant Dis. 106:2904–2910.

Schmitt, D. P., and Barker, K. R. 1981. Damage and Reproductive Potentials of *Pratylenchus brachyurus* and *Pratylenchus penetrans* on Soybean. J. Nematol. 13:327–332.

Schumacher, L. A., and Bird, G. W. 2025. Distribution of *Pratylenchus* spp. and *Heterodera glycines* in Michigan and Combined Reproduction on Peking and PI 437654 Soybean. Plant Health Prog. 10.1094/PHP-10-24-0101-RS

Schweiger, R., Heise, A. M., Persicke, M., and Muller, C. 2014. Interactions between the jasmonic and salicylic acid pathway modulate the plant metabolome and affect herbivores of different feeding types. Plant Cell Environ. 37:1574–1585.

Townshend, J. L. 1984. Anhydrobiosis in *Pratylenchus penetrans*. J. Nematol. 16:282–289.

Tylka, G. L., Gebhart, G. D., Marrett, C. C., and Mullaney, M. P. 2012. Evaluation of soybean varieties resistant to soybean cyst nematode in Iowa in 2012. Iowa State University Extension and Outreach. https://faculty.sites.iastate.edu/gltylka/files/inline-files/evaluation_of_scn-resistant_soybean_varieties_in_iowa_-_2018_ipm_52.pdf

Usovsky, M., Robert T. Robbins, Juliet Fultz Wilkes, Devany Crippen, Vijay Shankar, Tri D. Vuong, Paula Agudelo, and Nguyen, H. T. 2022. Classification methods and identification of reniform nematode resistance in known soybean cyst nematode resistant soybean genotypes. Plant Dis. 106:382–389.

Usovsky, M., Lakhssassi, N., Patil, G. B., Vuong, T. D., Piya, S., Hewezi, T., Robbins, R. T., Stupar, R. M., Meksem, K., and Nguyen, H. T. 2021. Dissecting nematode resistance regions in soybean revealed pleiotropic effect of soybean cyst and reniform nematode resistance genes. Plant Genome 14:e20083.

Usovsky, M., Gamage, V. A., Meinhardt, C. G., Dietz, N., Triller, M., Basnet, P., Gillman, J. D., Bilyeu, K. D., Song, Q., Dhital, B., Nguyen, A., Mitchum, M. G., and Scaboo, A. M. 2023. Loss-of-function of an alpha-SNAP gene confers resistance to soybean cyst nematode. Nat. Commun. 14:7629.

Verma, K., Kumari, K., Rawat, M., Devi, K., and Joshi, R. 2025. Crosstalk of Jasmonic acid and Salicylic acid with other Phytohormones Alleviates Abiotic and Biotic Stresses in Plants. J. Soil Sci. Plant Nutr. 25:4997–5019.

Yu, N., Lee, T. G., Rosa, D. P., Hudson, M., and Diers, B. W. 2016. Impact of *Rhg1* copy number, type, and interaction with *Rhg4* on resistance to Heterodera glycines in soybean. Theor. Appl. Genet. 129:2403–2412.

Zhao, M., Wu, S., Zhou, Q., Vivona, S., Cipriano, D. J., Cheng, Y., and Brunger, A. T. 2015. Mechanistic insights into the recycling machine of the SNARE complex. Nature 518:61–67.

Zunke, U. 1990. Observations on the Invasion and Endoparasitic Behavior of the Root Lesion Nematode *Pratylenchus-Penetrans*. J. Nematol. 22:309–320.

