## Supplementary material for "Soybean *Rhg1* resistance to *Heterodera glycines* hinders the parasitism of *Pratylenchus penetrans*": All supplemental information

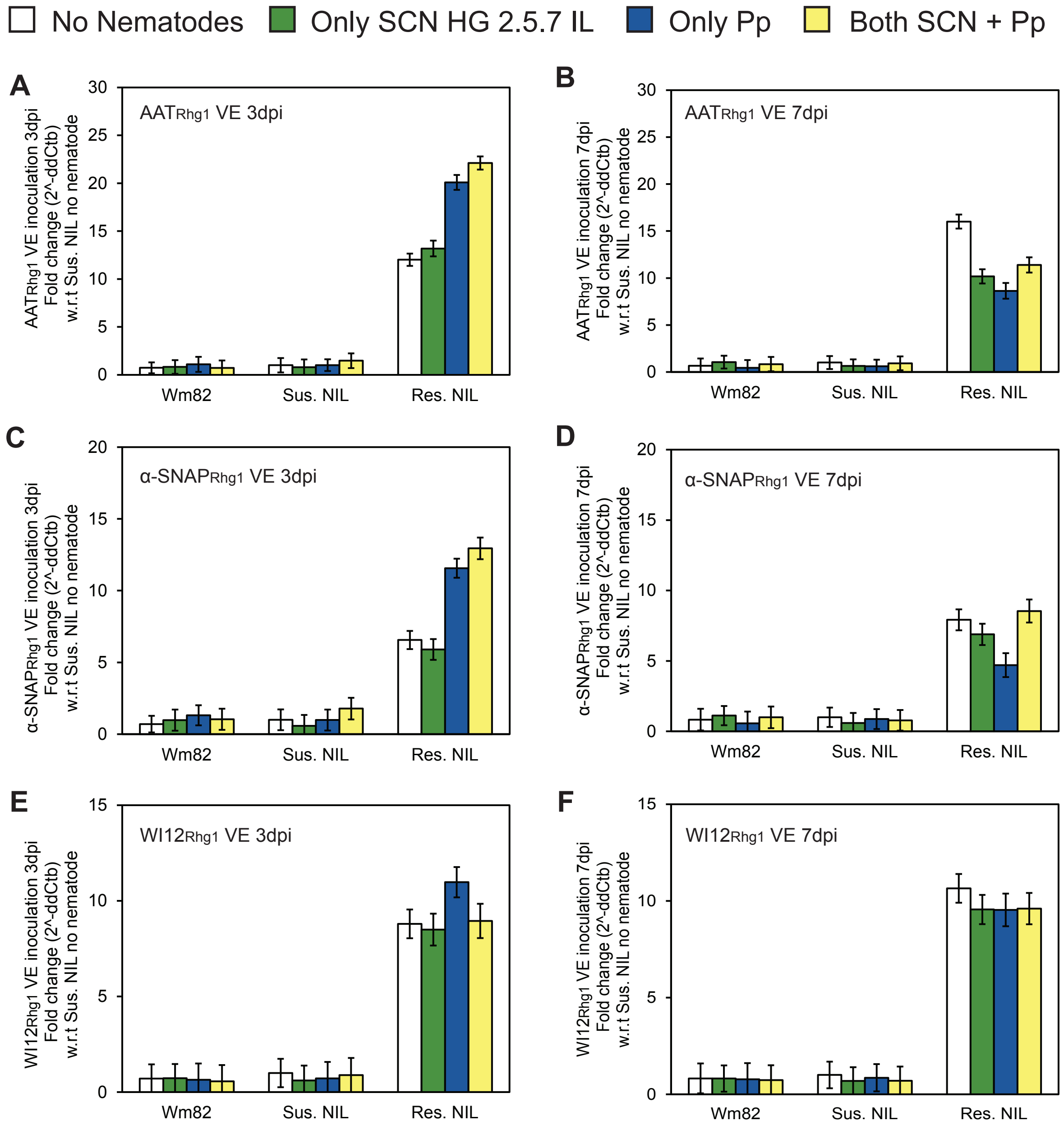

**Supplemental Figure 1. Same data as in Figure 4 but with data for all genotypes normalized to mean Sus. NIL no nematode control within same experiment, to show relative abundances between genotypes.** *Rhg1* transcript induction (fold change) compared to no-nematode controls within each plant genotype. The protein encoded by the analyzed transcript is noted in the upper-left corner of each graph. Wm82, Sus. NIL and Res. NIL plants were inoculated at VE stage and root samples from infection areas taken at: (A, C, E and G) 3 dpi, or (B, D, F and H) 7 dpi. No statistically significant differences between samples were observed using ANOVA and Tukey HSD test. The SCN HG 2.5.7 IL used partially overcomes *rhg1-b* resistance and represents the most prevalent HG type in the USA. Data are for two independent experiments with three replicate roots assessed within each experiment (n=6 for each bar).

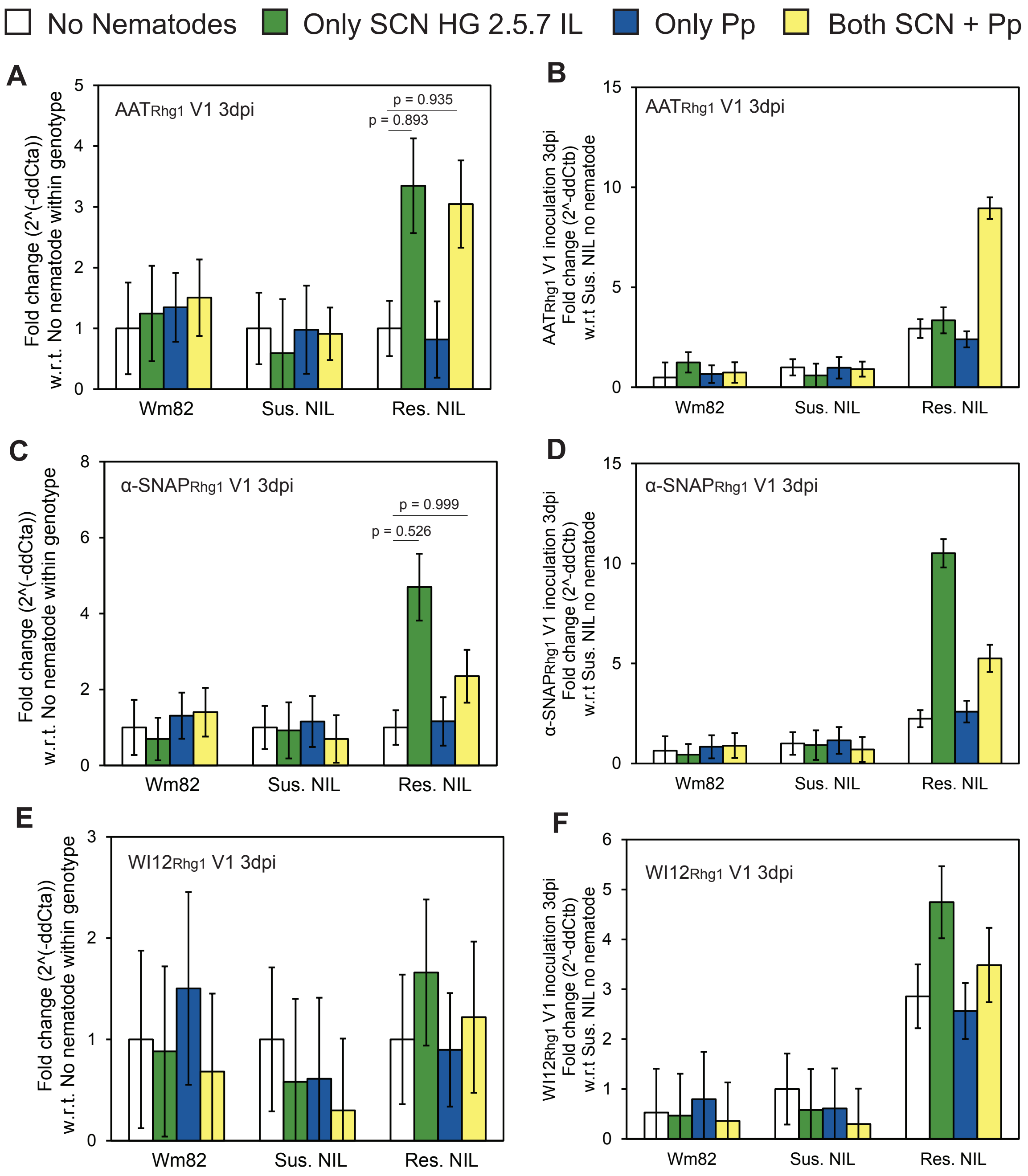

**Supplemental Figure 2. *Rhg1* transcript induction (fold change) for V1 plants at 3 dpi compared to no-nematode controls within each plant genotype.** The protein encoded by the analyzed transcript is noted in the upper-left corner of each graph. Wm82, Sus. NIL and Res. NIL plants were inoculated at V1 stage. (A, C, E) Data normalized to mean for no-nematode control of same genotype in same experiment. (B, D, F) Same data as A, C, E with data for all genotypes normalized to mean Sus. NIL no nematode control within same experiment, to show relative abundances between genotypes. Instances of possible significant differences of means are marked with p-values from ANOVA Tukey HSD tests ($p > 0.2$ if no p value is shown). The SCN HG 2.5.7 IL used partially overcomes *rhg1-b* resistance and represents the most prevalent HG type in the USA. All experiments were repeated twice with two replicate roots assessed within each experiment (n=4 for each bar).

**Supplementary Table 1.** List of all qPCR primers used in this study to quantify mRNA/transcript abundance

| Gene name | Primer Direction | 5' to 3' Primer Sequence | Reference |
|---|---|---|---|
| Glyma.18G022500 α-SNAP$_{Rhg1}$ | Forward | GCAATTGATGAAGAAGATGTTG | Haarith et al., 2025 |
| | Reverse | TCAAGTAATAGCCTCATGCTG | |
| Glyma.18G022700 WI12$_{Rhg1}$ | Forward | AGGTCACGTGTTGCCGTTG | Cook et al., 2012 |
| | Reverse | AAACCACACCAATAACAACAAAGCTCT | |
| Glyma.18G022400 AAT$_{Rhg1}$ | Forward | CGTGTAGAGTCCTTGAAGTACAGC | |
| | Reverse | ACCAGAGCTGTGATAGCCAACC | |
| Glyma.10G109500 AAT (JA) | Forward | GAGTTGATACCAGCCAACAG | Beyer et al., 2021 |
| | Reverse | GAGACCAACAAGCAGTGTAG | |
| Glyma.10G010100 NIMIN1 (SA) | Forward | ATGTTGAACACGGCATTCTC | |
| | Reverse | GGTACGGTGTGACTTTCTTG | |
| Glyma.11G079600 NREG1 (Reference gene) | Forward | TCCTGGCTGCTAACTACTTG | |
| | Reverse | GGTCTTGCGAATTTCCTCTG | |

**Supplementary Table 2.** ANOVA for Pp numbers in 1:1 SCN:Pp inoculation (Wm82, IL3045N and IL3849N)

Formula: aov(Sqrt(Pp count) ~ (Experiment/replicate)+Plant genotypes*Nematode inoculation)

| | Df | Sum. Sq | Mean Sq. | F value | Pr > F | Significance |
|---|---|---|---|---|---|---|
| **Experiment** | 1 | 4.476 | 4.4759 | 12.1744 | 0.001113 | ** |
| **Plant genotypes** | 2 | 33.379 | 16.6896 | 45.3959 | 2.011e-11 | *** |
| **Nematode inoculation** | 1 | 29.596 | 29.5964 | 80.5025 | 1.696e-11 | *** |
| **Experiment/replicate** | 8 | 9.173 | 1.1466 | 3.1189 | 0.007046 | ** |
| **Plant genotypes x Nematode inoculation** | 2 | 0.037 | 0.0185 | 0.0502 | 0.0951098 | |
| **Residuals** | 44 | 16.176 | 0.3676 | | | |
| **Significance : 0 '***' 0.001 '**' 0.01 '*'** | | | | | | |

| **Tukey HSD (p-values)** | | |
|---|---|---|
| Wm82+Pp | IL3849N+Pp | **0.0000004** |
| Wm82+Pp | IL3025N+Pp | **0.0000718** |
| Wm82+Both | Wm82+Pp | **0.0001694** |
| Wm82+Both | IL3849N+Pp | **0** |
| Wm82+Both | IL3849N+Both | **0.0000008** |
| Wm82+Both | IL3025N+Pp | **0** |
| Wm82+Both | IL3025N+Both | **0.0004221** |
| IL3849N+Pp | IL3025N+Pp | 0.6180058 |
| IL3849N+Both | Wm82+Pp | 0.5687882 |
| IL3849N+Both | IL3849N+Pp | **0.0000925** |
| IL3849N+Both | IL3025N+Pp | **0.0116435** |
| IL3849N+Both | IL3025N+Both | 0.3892712 |
| IL3025N+Both | Wm82+Pp | 0.9996929 |
| IL3025N+Both | IL3849N+Pp | **0.0000001** |
| IL3025N+Both | IL3025N+Pp | **0.0000274** |

**Supplementary Table 3.** ANOVA for SCN cyst numbers in 1:1 SCN:Pp inoculation (Wm82, IL3045N and IL3849N)

Formula: aov(log(Cyst count) ~ (Experiment/replicate)+Plant genotypes*Nematode inoculation)

| | Df | Sum. Sq | Mean Sq. | F value | Pr > F | Significance |
|---|---|---|---|---|---|---|
| **Experiment** | 1 | 14.217 | 14.217 | 52.4515 | 4.367e-09 | *** |
| **Plant genotypes** | 2 | 92.931 | 46.465 | 171.7184 | <2.2e-16 | *** |
| **Nematode inoculation** | 1 | 0.001 | 0.001 | 0.0045 | 0.94687 | |
| **Experiment/replicate** | 8 | 1.438 | 0.180 | 0.6641 | 0.71986 | |
| **Plant genotypes x Nematode inoculation** | 2 | 2.516 | 1.258 | 4.6485 | 0.01462 | * |
| **Residuals** | 45 | 12.177 | 0.271 | | | |
| **Significance : 0 ‘***’ 0.001 ‘**’ 0.01 ‘*’** | | | | | | |

| **Tukey HSD (p-values)** | | |
|---|---|---|
| Wm82+SCN | IL3849N+SCN | **0.000000** |
| Wm82+SCN | IL3025N+SCN | **0.003122** |
| Wm82+Both | Wm82+SCN | 0.168464 |
| Wm82+Both | IL3849N+SCN | **0.000000** |
| Wm82+Both | IL3849N+Both | **0.000000** |
| Wm82+Both | IL3025N+SCN | 0.628442 |
| Wm82+Both | IL3025N+Both | 0.912637 |
| IL3849N+SCN | IL3025N+SCN | **0.000000** |
| IL3849N+Both | Wm82+SCN | **0.000000** |
| IL3849N+Both | IL3849N+SCN | 0.500285 |
| IL3849N+Both | IL3025N+SCN | **0.000000** |
| IL3849N+Both | IL3025N+Both | **0.000000** |
| IL3025N+Both | Wm82+Pp | 0.050215 |
| IL3025N+Both | IL3849N+Pp | **0** |
| IL3025N+Both | IL3025N+Pp | **0.0002341** |

**Supplementary Table 4.** ANOVA for Pp numbers in 2:1 SCN:Pp inoculation(Wm82, IL3045N and IL3849N)

Formula: aov(log(Pp count) ~ (Experiment/replicate)+Plant genotypes*Nematode inoculation)

| | Df | Sum. Sq | Mean Sq. | F value | Pr > F | Significance |
|---|---|---|---|---|---|---|
| **Experiment** | 1 | 0.8422 | 0.8422 | 5.9161 | 0.01904 | * |
| **Plant genotypes** | 2 | 10.9284 | 5.4642 | 38.3814 | 1.876e-10 | *** |
| **Nematode inoculation** | 1 | 7.9374 | 7.9374 | 55.7533 | 2.081e-09 | *** |
| **Experiment/replicate** | 8 | 0.4230 | 0.0529 | 0.3714 | 0.93021 | |
| **Plant genotypes x Nematode inoculation** | 2 | 0.1748 | 0.0874 | 0.6140 | 0.54565 | |
| **Residuals** | 45 | 6.4065 | 0.1424 | | | |
| **Significance : 0 '***' 0.001 '**' 0.01 '*'** | | | | | | |

| **Tukey HSD (p-values)** | | |
|---|---|---|
| Wm82+Pp | IL3849N+Pp | **0.0000364** |
| Wm82+Pp | IL3025N+Pp | 0.4505769 |
| Wm82+Both | Wm82+Pp | **0.0003304** |
| Wm82+Both | IL3025N+Pp | **0.0000007** |
| Wm82+Both | IL3849N+Pp | **0** |
| Wm82+Both | IL3849N+Both | **0.0000004** |
| Wm82+Both | IL3025N+Both | 0.5168828 |
| IL3849N+Pp | IL3025N+Pp | **0.0114356** |
| IL3849N+Both | Wm82+Pp | 0.3737783 |
| IL3849N+Both | IL3849N+Pp | **0.0163912** |
| IL3849N+Both | IL3025N+Pp | 0.9999942 |
| IL3849N+Both | IL3025N+Both | **0.0001528** |
| IL3025N+Both | Wm82+Pp | 0.050215 |
| IL3025N+Both | IL3849N+Pp | **0** |
| IL3025N+Both | IL3025N+Pp | **0.0002341** |

**Supplementary Table 5.** ANOVA for SCN cyst numbers in 2:1 SCN:Pp inoculation (Wm82, IL3045N and IL3849N)

Formula: aov(Sqrt(Cyst count) ~ (Experiment/replicate)+Plant genotypes*Nematode inoculation)

| | Df | Sum. Sq | Mean Sq. | F value | Pr > F | Significance |
|---|---|---|---|---|---|---|
| **Experiment** | 1 | 221.4 | 221.44 | 21.4969 | 3.053e-05 | *** |
| **Plant genotypes** | 2 | 5954.2 | 2977.08 | 289.0036 | <2.2e-16 | *** |
| **Nematode inoculation** | 1 | 2.4 | 2.42 | 0.2350 | 0.6301709 | |
| **Experiment/replicate** | 8 | 404.6 | 50.58 | 4.9097 | 0.0002125 | *** |
| **Plant genotypes x Nematode inoculation** | 2 | 5.0 | 2.50 | 0.2430 | 0.7852932 | |
| **Residuals** | 45 | 463.6 | 10.30 | | | |
| **Significance : 0 '***' 0.001 '**' 0.01 '*'** | | | | | | |

| **Tukey HSD (p-values)** | | |
|---|---|---|
| Wm82+SCN | IL3849N+SCN | **0.0000000** |
| Wm82+SCN | IL3025N+SCN | **0.0000632** |
| Wm82+Both | Wm82+SCN | 1.0000000 |
| Wm82+Both | IL3849N+SCN | **0.0000000** |
| Wm82+Both | IL3849N+Both | 0.0000000 |
| Wm82+Both | IL3025N+SCN | **0.0000668** |
| Wm82+Both | IL3025N+Both | **0.0010353** |
| IL3849N+SCN | IL3025N+SCN | **0.0000000** |
| IL3849N+Both | Wm82+SCN | 0.0000000 |
| IL3849N+Both | IL3849N+SCN | 1.0000000 |
| IL3849N+Both | IL3025N+SCN | **0.0000000** |
| IL3849N+Both | IL3025N+Both | 0.0000000 |
| IL3025N+Both | Wm82+SCN | **0.0009817** |
| IL3025N+Both | IL3849N+SCN | **0.0000000** |
| IL3025N+Both | IL3025N+SCN | 0.9563879 |

**Supplementary Table 6.** ANOVA for Total Pp numbers on NILs

Formula: aov(sqrt(Pp count) ~ (Experiment/replicate)+Plant genotypes*Nematode inoculation)

| | **Df** | **Sum. Sq** | **Mean Sq.** | **F value** | **Pr > F** | **Significance** |
|---|---|---|---|---|---|---|
| **Experiment** | 1 | 103.184 | 103.184 | 122.6661 | 1.926e-14 | *** |
| **Plant genotypes** | 2 | 26.913 | 13.456 | 15.9970 | 5.649e-06 | *** |
| **Nematode inoculation** | 1 | 14.541 | 14.541 | 17.2866 | 0.0001423 | *** |
| **Experiment/replicate** | 8 | 5.670 | 0.709 | 0.8426 | 0.5706470 | |
| **Plant genotypes x Nematode inoculation** | 2 | 4.094 | 2.047 | 2.4333 | 0.0992143 | |
| **Residuals** | 45 | 37.853 | 0.841 | | | |
| **Significance : 0 '***' 0.001 '**' 0.01 '*'** | | | | | | |

| **Tukey HSD (p-values)** | | |
|---|---|---|
| Wm82+Pp | Wm82+Both | **0.0018532** |
| Wm82+Pp | Sus. NIL+Pp | 0.9932719 |
| Wm82+Pp | Sus. NIL+Both | 0.5104347 |
| Wm82+Pp | Res. NIL+Pp | 0.0994669 |
| Wm82+Pp | Res. NIL+Both | 0.9609094 |
| Wm82+Both | Sus. NIL+Both | 0.172806 |
| Wm82+Both | Res. NIL+Both | **0.0001337** |
| Sus. NIL+Pp | Wm82+Both | **0.0095477** |
| Sus. NIL+Pp | Sus. NIL+Both | 0.8400984 |
| Sus. NIL+Pp | Res. NIL+Pp | **0.0258896** |
| Sus. NIL+Pp | Res. NIL+Both | 0.7362778 |
| Sus. NIL+Both | Res. NIL+Both | 0.1255234 |
| Res. NIL+Pp | Wm82+Both | **0.0000003** |
| Res. NIL+Pp | Sus. NIL+Both | **0.0008159** |
| Res. NIL+Pp | Res. NIL+Both | 0.4436716 |

**Supplementary Table 7.** ANOVA for Total Pp Sand:Root ratio on NILs

Formula: aov(sqrt(Pp Sand:Root count ratio) ~ (Experiment/replicate)+Plant genotypes*Nematode inoculation)

| | Df | Sum. Sq | Mean Sq. | F value | Pr > F | Significance |
|---|---|---|---|---|---|---|
| **Experiment** | 1 | 2.5698 | 2.5698 | 8.2737 | 0.006128 | ** |
| **Plant genotypes** | 2 | 0.7019 | 0.3509 | 1.1298 | 0.332080 | |
| **Nematode inoculation** | 1 | 9.8983 | 9.8983 | 31.8680 | 1.047e-06 | *** |
| **Experiment/replicate** | 8 | 2.2440 | 0.2805 | 0.9031 | 0.522299 | |
| **Plant genotypes x Nematode inoculation** | 2 | 7.9391 | 3.9695 | 12.7801 | 4.024e-05 | *** |
| **Residuals** | 45 | 13.9771 | 0.3106 | | | |
| **Significance : 0 '***' 0.001 '**' 0.01 '*'** | | | | | | |

| **Tukey HSD (p-values)** | | |
|---|---|---|
| Wm82+Pp | Wm82+Both | 0.9992374 |
| Wm82+Pp | Sus. NIL+Pp | 0.5181065 |
| Wm82+Pp | Sus. NIL+Both | 0.9977317 |
| Wm82+Pp | Res. NIL+Pp | **0.0007679** |
| Wm82+Pp | Res. NIL+Both | 0.0747088 |
| Wm82+Both | Sus. NIL+Both | 0.9999991 |
| Wm82+Both | Res. NIL+Both | 0.1598771 |
| Sus. NIL+Pp | Wm82+Both | 0.3133064 |
| Sus. NIL+Pp | Sus. NIL+Both | 0.2696013 |
| Sus. NIL+Pp | Res. NIL+Pp | 0.0927678 |
| Sus. NIL+Pp | Res. NIL+Both | **0.0005679** |
| Sus. NIL+Both | Res. NIL+Both | 0.90591 |
| Res. NIL+Pp | Wm82+Both | **0.0002492** |
| Res. NIL+Pp | Sus. NIL+Both | **0.0001862** |
| Res. NIL+Pp | Res. NIL+Both | **0.0000001** |

**Supplementary Table 8.** ANOVA for Total Pp Female:Male ratio on NILs

aov(sqrt(Female:Male count ratio) ~ (Experiment/replicate)+Plant genotypes*Nematode inoculation)

| | Df | Sum. Sq | Mean Sq. | F value | Pr > F | Significance |
|---|---|---|---|---|---|---|
| **Experiment** | 1 | 0.2124 | 0.21241 | 1.4121 | 0.2402 | |
| **Plant genotypes** | 2 | 4.7374 | 2.36870 | 15.7469 | 4.724e-06 | *** |
| **Nematode inoculation** | 1 | 0.3330 | 0.33301 | 2.2139 | 0.1429 | |
| **Experiment/replicate** | 2 | 0.1788 | 0.08940 | 0.5943 | 0.5557 | |
| **Plant genotypes x Nematode inoculation** | 2 | 0.1189 | 0.05945 | 0.3952 | 0.6756 | |
| **Residuals** | 51 | 7.6716 | 0.15042 | | | |
| **Significance : 0 '***' 0.001 '**' 0.01 '*'** | | | | | | |

| **Tukey HSD (p-value)** | | |
|---|---|---|
| Wm82+Pp | Wm82+Both | 0.9589550 |
| Wm82+Pp | Sus. NIL+Pp | 0.9999986 |
| Wm82+Pp | Sus. NIL+Both | 0.8754047 |
| Wm82+Pp | Res. NIL+Pp | **0.0286405** |
| Wm82+Pp | Res. NIL+Both | 0.0712761 |
| Wm82+Both | Sus. NIL+Both | 0.9997857 |
| Wm82+Both | Res. NIL+Both | **0.0080514** |
| Sus. NIL+Pp | Wm82+Both | 0.9759452 |
| Sus. NIL+Pp | Sus. NIL+Both | 0.9121446 |
| Sus. NIL+Pp | Res. NIL+Pp | **0.0221532** |
| Sus. NIL+Pp | Res. NIL+Both | 0.0565343 |
| Sus. NIL+Both | Res. NIL+Both | **0.0036522** |
| Res. NIL+Pp | Wm82+Both | **0.0027149** |
| Res. NIL+Pp | Sus. NIL+Both | **0.0011828** |
| Res. NIL+Pp | Res. NIL+Both | 0.9990226 |

**Supplementary Table 9.** ANOVA for Total Pp Juveniles (J2+J3+J4) on NILs

| | Df | Sum. Sq | Mean Sq. | F value | Pr > F | Significance |
|---|---|---|---|---|---|---|
| **Experiment** | 1 | 49.310 | 49.310 | 103.5823 | 2.988e-13 | *** |
| **Plant genotypes** | 2 | 23.238 | 11.619 | 24.4071 | 6.626e-08 | *** |
| **Nematode inoculation** | 1 | 11.532 | 11.532 | 24.2241 | 1.193e-05 | *** |
| **Experiment/replicate** | 8 | 4.080 | 0.510 | 1.0712 | 0.39996 | |
| **Plant genotypes x Nematode inoculation** | 2 | 4.297 | 2.149 | 4.5136 | 0.01635 | * |
| **Residuals** | 45 | 21.422 | 0.476 | | | |
| **Significance : 0 '***' 0.001 '**' 0.01 '*'** | | | | | | |

| **Tukey HSD (p-value)** | | |
|---|---|---|
| Wm82+Pp | Wm82+Both | **0.0000487** |
| Wm82+Pp | Sus. NIL+Pp | 0.8636626 |
| Wm82+Pp | Sus. NIL+Both | 0.1015930 |
| Wm82+Pp | Res. NIL+Pp | 0.0752190 |
| Wm82+Pp | Res. NIL+Both | 0.8762907 |
| Wm82+Both | Sus. NIL+Both | 0.1126620 |
| Wm82+Both | Res. NIL+Both | **0.0000012** |
| Sus. NIL+Pp | Wm82+Both | **0.0018727** |
| Sus. NIL+Pp | Sus. NIL+Both | 0.6445899 |
| Sus. NIL+Pp | Res. NIL+Pp | **0.0036033** |
| Sus. NIL+Pp | Res. NIL+Both | 0.2393802 |
| Sus. NIL+Both | Res. NIL+Both | **0.0058241** |
| Res. NIL+Pp | Wm82+Both | **0.0000000** |
| Res. NIL+Pp | Sus. NIL+Both | **0.0000000** |
| Res. NIL+Pp | Res. NIL+Both | 0.5404546 |

**Supplementary Table 10.** ANOVA for Total Pp Juveniles (J2+J3+J4):Adults (Male+Female) on NILs

| | Df | Sum. Sq | Mean Sq. | F value | Pr > F | Significance |
|---|---|---|---|---|---|---|
| **Experiment** | 1 | 0.1004 | 0.10043 | 1.1352 | 0.292359 | |
| **Plant genotypes** | 2 | 1.4132 | 0.70662 | 7.9874 | 0.001075 | ** |
| **Nematode inoculation** | 1 | 0.4680 | 0.46803 | 5.2903 | 0.026135 | * |
| **Experiment/replicate** | 8 | 0.7269 | 0.09086 | 1.0271 | 0.430151 | |
| **Plant genotypes x Nematode inoculation** | 2 | 0.3017 | 0.15083 | 1.7049 | 0.193321 | |
| **Residuals** | 51 | 3.9810 | 0.08847 | | | |
| **Significance : 0 '***' 0.001 '**' 0.01 '*'** | | | | | | |

| **Tukey HSD (p-value)** | | |
|---|---|---|
| Wm82+Pp | Wm82+Both | 0.1379612 |
| Wm82+Pp | Sus. NIL+Pp | 0.7765115 |
| Wm82+Pp | Sus. NIL+Both | 0.0702945 |
| Wm82+Pp | Res. NIL+Pp | 0.9867943 |
| Wm82+Pp | Res. NIL+Both | 0.9799576 |
| Wm82+Both | Sus. NIL+Both | 0.9996147 |
| Wm82+Both | Res. NIL+Both | **0.0259743** |
| Sus. NIL+Pp | Wm82+Both | 0.827706 |
| Sus. NIL+Pp | Sus. NIL+Both | 0.653358 |
| Sus. NIL+Pp | Res. NIL+Pp | 0.3818849 |
| Sus. NIL+Pp | Res. NIL+Both | 0.3468304 |
| Sus. NIL+Both | Res. NIL+Both | **0.0113626** |
| Res. NIL+Pp | Wm82+Both | **0.0305982** |
| Res. NIL+Pp | Sus. NIL+Both | **0.013538** |
| Res. NIL+Pp | Res. NIL+Both | 0.9999998 |

**Supplementary Table 11.** ANOVA for Total SCN cysts on NILs

| | Df | Sum. Sq | Mean Sq. | F value | Pr > F | Significance |
|---|---|---|---|---|---|---|
| **Experiment** | 1 | 13.0096 | 13.0096 | 115.6445 | 5.082e-14 | *** |
| **Plant genotypes** | 2 | 1.0089 | 0.5045 | 4.4843 | 0.01675 | * |
| **Nematode inoculation** | 1 | 2.4579 | 2.4579 | 21.8488 | 2.698e-05 | *** |
| **Experiment/replicate** | 8 | 0.2749 | 0.0344 | 0.3054 | 0.96010 | |
| **Plant genotypes x Nematode inoculation** | 2 | 0.3164 | 0.1582 | 1.4064 | 0.25560 | |
| **Residuals** | 45 | 5.0624 | 0.1125 | | | |
| **Significance : 0 '***' 0.001 '**' 0.01 '*'** | | | | | | |

| **Tukey HSD (p-values)** | | |
|---|---|---|
| Wm82+SCN | Wm82+Both | **0.0039955** |
| Wm82+SCN | Sus. NIL+SCN | 0.9978039 |
| Wm82+SCN | Sus. NIL+Both | **0.0371747** |
| Wm82+SCN | Res. NIL+SCN | 0.0572952 |
| Wm82+SCN | Res. NIL+Both | **0.0007447** |
| Wm82+Both | Sus. NIL+Both | 0.9643226 |
| Wm82+Both | Res. NIL+Both | 0.9937006 |
| Sus. NIL+SCN | Wm82+Both | **0.0139943** |
| Sus. NIL+SCN | Sus. NIL+Both | 0.1046342 |
| Sus. NIL+SCN | Res. NIL+SCN | 0.1516324 |
| Sus. NIL+SCN | Res. NIL+Both | **0.0028780** |
| Sus. NIL+Both | Res. NIL+Both | 0.7510163 |
| Res. NIL+SCN | Wm82+Both | 0.9202250 |
| Res. NIL+SCN | Sus. NIL+Both | 0.9999748 |
| Res. NIL+SCN | Res. NIL+Both | 0.6445139 |

**Supplementary Table 12.** ANOVA for RT q-PCR transcript abundance for 3dpi VE inoculation 1:1 SCN:Pp. Analyses were done on ddCt values = (Ct sample – CtNREG1) - avg(Ct No nematode sample of the same plant genotype – CtNREG1).

**$AAT_{Rhg1}$ 3dpi VE inoculation**

| | Df | Sum. Sq | Mean Sq. | F value | Pr > F | Significance |
|---|---|---|---|---|---|---|
| **Experiment** | 1 | 5.456 | 5.4560 | 4.8545 | 0.03178 | * |
| **Plant x Nematode Treatment** | 11 | 9.183 | 0.8348 | 0.7428 | 0.69323 | |
| **Experiment/replicate (root)** | 4 | 41.001 | 10.2504 | 9.1203 | 1.003e-05 | *** |
| **Residuals** | 55 | 61.815 | 1.1239 | | | |
| **Significance : 0 '***' 0.001 '**' 0.01 '*'** | | | | | | |

**$\alpha$-$SNAP_{Rhg1}$ 3dpi VE inoculation**

| | Df | Sum. Sq | Mean Sq. | F value | Pr > F | Significance |
|---|---|---|---|---|---|---|
| **Experiment** | 1 | 6.559 | 6.5590 | 5.7852 | 0.01955 | * |
| **Plant x Nematode Treatment** | 11 | 19.880 | 1.8073 | 1.5941 | 0.12644 | |
| **Experiment/replicate (root)** | 4 | 47.520 | 11.8800 | 10.4784 | 2.212e-06 | *** |
| **Residuals** | 55 | 62.357 | 1.1338 | | | |
| **Significance : 0 '***' 0.001 '**' 0.01 '*'** | | | | | | |

**$WI12_{Rhg1}$ 3dpi VE inoculation**

| | Df | Sum. Sq | Mean Sq. | F value | Pr > F | Significance |
|---|---|---|---|---|---|---|
| **Experiment** | 1 | 0.9014 | 0.9014 | 2.1180 | 0.1513 | |
| **Plant x Nematode Treatment** | 11 | 4.8492 | 0.4408 | 1.0358 | 0.4289 | |
| **Experiment/replicate (root)** | 4 | 24.2835 | 6.0709 | 14.2642 | 4.749e-08 | *** |
| **Residuals** | 55 | 23.4081 | 0.4256 | | | |
| **Significance : 0 '***' 0.001 '**' 0.01 '*'** | | | | | | |

**$\alpha$-$SNAP_{Rhg1}HC$ 3dpi VE inoculation**

| | Df | Sum. Sq | Mean Sq. | F value | Pr > F | Significance |
|---|---|---|---|---|---|---|
| **Experiment** | 1 | 1.0645 | 1.0645 | 5.7852 | 0.01955 | * |
| **Plant x Nematode Treatment** | 3 | 0.7744 | 0.7744 | 1.5941 | 0.12644 | |
| **Experiment/replicate (root)** | 4 | 6.6203 | 6.6203 | 10.4784 | 2.212e-06 | *** |
| **Residuals** | 15 | 0.2607 | 0.2607 | | | |
| **Significance : 0 '***' 0.001 '**' 0.01 '*'** | | | | | | |

| **AAT (JA response) 3dpi VE inoculation** | | | | | | |
|---|---|---|---|---|---|---|
| | Df | Sum. Sq | Mean Sq. | F value | Pr > F | Significance |
| **Experiment** | 1 | 0.210 | 0.2097 | 0.3154 | 0.5767 | |
| **Plant x Nematode Treatment** | 11 | 5.761 | 0.5238 | 0.7876 | 0.6511 | |
| **Experiment/replicate (root)** | 4 | 62.289 | 15.5722 | 23.4174 | 2.438e-11 | *** |
| **Residuals** | 55 | 36.574 | 0.6650 | | | |
| **Significance : 0 ‘***’ 0.001 ‘**’ 0.01 ‘*’** | | | | | | |

| **NIMIN1 (SA response) 3dpi VE inoculation** | | | | | | |
|---|---|---|---|---|---|---|
| | Df | Sum. Sq | Mean Sq. | F value | Pr > F | Significance |
| **Experiment** | 1 | 19.621 | 19.6207 | 28.1187 | 2.087e-06 | *** |
| **Plant x Nematode Treatment** | 11 | 53.984 | 4.9076 | 7.0331 | 2.848e-07 | *** |
| **Experiment/replicate (root)** | 4 | 37.738 | 9.4345 | 13.5207 | 9.706e-08 | *** |
| **Residuals** | 55 | 38.378 | 0.6978 | | | |
| **Significance : 0 ‘***’ 0.001 ‘**’ 0.01 ‘*’** | | | | | | |

**Supplementary Table 13.** ANOVA for RT q-PCR transcript abundance for 7dpi VE inoculation 1:1 SCN:Pp. Analyses were done on ddCt values = (Ct sample – CtNREG1) - avg(Ct No nematode sample of the same plant genotype – CtNREG1).

**$AAT_{Rhg1}$ 3dpi VE inoculation**

| | Df | Sum. Sq | Mean Sq. | F value | Pr > F | Significance |
|---|---|---|---|---|---|---|
| **Experiment** | 1 | 1.711 | 1.7713 | 1.8180 | 0.1830820 | |
| **Plant x Nematode Treatment** | 11 | 4.747 | 0.4316 | 0.4585 | 0.9205702 | |
| **Experiment/replicate (root)** | 4 | 25.400 | 6.3499 | 6.7458 | 0.0001717 | *** |
| **Residuals** | 55 | 51.772 | 0.9413 | | | |
| **Significance : 0 '***' 0.001 '**' 0.01 '*'** | | | | | | |

**α-$SNAP_{Rhg1}$ 3dpi VE inoculation**

| | Df | Sum. Sq | Mean Sq. | F value | Pr > F | Significance |
|---|---|---|---|---|---|---|
| **Experiment** | 1 | 0.5753 | 0.57525 | 1.0549 | 0.30889 | |
| **Plant x Nematode Treatment** | 11 | 4.2437 | 0.38579 | 0.7074 | 0.72612 | |
| **Experiment/replicate (root)** | 4 | 6.7825 | 1.69564 | 3.1093 | 0.02231 | * |
| **Residuals** | 55 | 29.9937 | 0.54534 | | | |
| **Significance : 0 '***' 0.001 '**' 0.01 '*'** | | | | | | |

**$WI12_{Rhg1}$ 3dpi VE inoculation**

| | Df | Sum. Sq | Mean Sq. | F value | Pr > F | Significance |
|---|---|---|---|---|---|---|
| **Experiment** | 1 | 0.0001 | 0.00007 | 0.0001 | 0.991382 | |
| **Plant x Nematode Treatment** | 11 | 2.2066 | 0.20060 | 0.3510 | 0.969036 | |
| **Experiment/replicate (root)** | 4 | 10.4876 | 2.62191 | 4.5881 | 0.002868 | ** |
| **Residuals** | 55 | 31.4299 | 0.57145 | | | |
| **Significance : 0 '***' 0.001 '**' 0.01 '*'** | | | | | | |

**α-$SNAP_{Rhg1}$HC 3dpi VE inoculation**

| | Df | Sum. Sq | Mean Sq. | F value | Pr > F | Significance |
|---|---|---|---|---|---|---|
| **Experiment** | 1 | 0.154 | 0.1535 | 0.4299 | 0.5220 | |
| **Plant x Nematode Treatment** | 3 | 0.261 | 0.0871 | 0.2440 | 0.8643 | |
| **Experiment/replicate (root)** | 4 | 92.762 | 23.1906 | 64.9315 | 2.735e-09 | *** |
| **Residuals** | 15 | 5.357 | 0.3572 | | | |
| **Significance : 0 '***' 0.001 '**' 0.01 '*'** | | | | | | |

### AAT (JA response) 3dpi VE inoculation

| | Df | Sum. Sq | Mean Sq. | F value | Pr > F | Significance |
|---|---|---|---|---|---|---|
| **Experiment** | 1 | 9.498 | 9.498 | 19.3388 | 5.061e-05 | *** |
| **Plant x Nematode Treatment** | 11 | 4.119 | 0.374 | 0.7625 | 0.6747 | |
| **Experiment/replicate (root)** | 4 | 170.788 | 42.697 | 86.9339 | <2.2e-16 | *** |
| **Residuals** | 55 | 27.013 | 0.491 | | | |
| **Significance : 0 '***' 0.001 '**' 0.01 '*'** | | | | | | |

### NIMIN1 (SA response) 3dpi VE inoculation

| | Df | Sum. Sq | Mean Sq. | F value | Pr > F | Significance |
|---|---|---|---|---|---|---|
| **Experiment** | 1 | 7.284 | 7.284 | 7.0647 | 0.01027 | * |
| **Plant x Nematode Treatment** | 11 | 16.375 | 1.489 | 1.4438 | 0.18027 | |
| **Experiment/replicate (root)** | 4 | 151.174 | 37.793 | 36.6539 | 6.423e-15 | *** |
| **Residuals** | 55 | 56.710 | 1.031 | | | |
| **Significance : 0 '***' 0.001 '**' 0.01 '*'** | | | | | | |

**Supplementary Table 14.** ANOVA for RT q-PCR transcript abundance for 3dpi V1 inoculation 1:1 SCN:Pp. Analyses were done on ddCt values = (Ct sample – CtNREG1) - avg(Ct No nematode sample of the same plant genotype – CtNREG1).

### $AAT_{Rhg1}$ 3dpi VE inoculation

| | Df | Sum. Sq | Mean Sq. | F value | Pr > F | Significance |
|---|---|---|---|---|---|---|
| **Experiment** | 1 | 11.146 | 11.1458 | 4.7505 | 0.03653 | * |
| **Plant x Nematode Treatment** | 11 | 23.710 | 2.1554 | 0.9187 | 0.53433 | |
| **Experiment/replicate (root)** | 2 | 4.453 | 2.2265 | 0.9490 | 0.39745 | |
| **Residuals** | 33 | 77.425 | 2.3462 | | | |
| **Significance : 0 '***' 0.001 '**' 0.01 '*'** | | | | | | |

### α-$SNAP_{Rhg1}$ 3dpi VE inoculation

| | Df | Sum. Sq | Mean Sq. | F value | Pr > F | Significance |
|---|---|---|---|---|---|---|
| **Experiment** | 1 | 10.949 | 10.9491 | 5.5854 | 0.02415 | * |
| **Plant x Nematode Treatment** | 11 | 25.812 | 2.3466 | 1.1970 | 0.32688 | |
| **Experiment/replicate (root)** | 2 | 12.521 | 6.2607 | 3.1938 | 0.05396 | * |
| **Residuals** | 33 | 64.690 | 1.9603 | | | |
| **Significance : 0 '***' 0.001 '**' 0.01 '*'** | | | | | | |

### $WI12_{Rhg1}$ 3dpi VE inoculation

| | Df | Sum. Sq | Mean Sq. | F value | Pr > F | Significance |
|---|---|---|---|---|---|---|
| **Experiment** | 1 | 1.150 | 1.1501 | 1.0715 | 0.3081 | |
| **Plant x Nematode Treatment** | 11 | 19.808 | 1.8008 | 1.6777 | 0.1227 | |
| **Experiment/replicate (root)** | 2 | 2.368 | 1.1840 | 1.1031 | 0.3438 | |
| **Residuals** | 33 | 35.420 | 1.0733 | | | |
| **Significance : 0 '***' 0.001 '**' 0.01 '*'** | | | | | | |

### α-$SNAP_{Rhg1}$HC 3dpi VE inoculation

| | Df | Sum. Sq | Mean Sq. | F value | Pr > F | Significance |
|---|---|---|---|---|---|---|
| **Experiment** | 1 | 1.69 | 1.69 | 0.0175 | 0.8963569 | |
| **Plant x Nematode Treatment** | 11 | 70.75 | 6.43 | 0.0667 | 0.9999709 | |
| **Experiment/replicate (root)** | 2 | 2834.42 | 1417.21 | 14.6904 | 0.0001972 | *** |
| **Residuals** | 17 | 1640.03 | 96.47 | | | |
| **Significance : 0 '***' 0.001 '**' 0.01 '*'** | | | | | | |

| AAT (JA response) 3dpi VE inoculation | | | | | | |
|---|---|---|---|---|---|---|
| | Df | Sum. Sq | Mean Sq. | F value | Pr > F | Significance |
| **Experiment** | 1 | 1.272 | 1.2724 | 0.6445 | 0.42785 | |
| **Plant x Nematode Treatment** | 11 | 24.857 | 2.2598 | 1.1445 | 0.36080 | |
| **Experiment/replicate (root)** | 2 | 15.736 | 7.8679 | 3.9850 | 0.02817 | * |
| **Residuals** | 33 | 65.154 | 1.9744 | | | |
| **Significance : 0 '***' 0.001 '**' 0.01 '*'** | | | | | | |

| NIMIN1 (SA response) 3dpi VE inoculation | | | | | | |
|---|---|---|---|---|---|---|
| | Df | Sum. Sq | Mean Sq. | F value | Pr > F | Significance |
| **Experiment** | 1 | 0.015 | 0.0152 | 0.0080 | 0.929478 | |
| **Plant x Nematode Treatment** | 11 | 74.317 | 6.7561 | 3.5360 | 0.002533 | ** |
| **Experiment/replicate (root)** | 2 | 14.637 | 7.3183 | 3.8303 | 0.032259 | * |
| **Residuals** | 32 | 61.140 | 1.9106 | | | |
| **Significance : 0 '***' 0.001 '**' 0.01 '*'** | | | | | | |
